# Integrative genomic study of mutation dynamics and Evolutionary trends in SARS-CoV-2 omicron BA.3

**DOI:** 10.1101/2025.08.03.668320

**Authors:** Milton S Kambarami

## Abstract

This study investigates the evolutionary dynamics of the SARS-CoV-2 BA.3 lineage with an emphasis on saltation-driven adaptation. Using 81 high-coverage BA.3 genome sequences and 1,011 complete patient status records obtained from GISAID, we conducted a comprehensive analysis that integrated temporal, mutational, phylogenetic, and selection pressure assessments. Temporal analysis of patient records revealed that BA.3 sequences were predominantly collected between November 2021 and March 2022, with highly mutated variants emerging in the last quarter of 2024. Pairwise alignment of the spike gene demonstrated near-identical sequences among recent isolates from Gauteng Province and subtle yet significant differences in a KwaZulu Natal isolate when compared to the Wuhan Hu-1 reference strain. Domain-specific mutation mapping showed that mutations were concentrated in key functional regions of the spike protein, particularly within the receptor-binding domain and its binding motif. Phylogenetic reconstruction and pervasive selection analysis further revealed that recent BA.3 variants form a divergent clade characterized by extensive adaptive mutations. These findings indicate that saltatory events play a critical role in shaping the genetic landscape of BA.3, with important implications for viral infectivity, immune escape, and the design of next-generation vaccines.

## Introduction

The SARS-CoV-2 pandemic has been marked by the rapid emergence and global spread of multiple viral variants. Among these, the Omicron variant is distinguished by an unusually high number of mutations in its spike protein, which has given rise to several sublineages such as BA.1, BA.2, and BA.3 (Desingu, Nagarajan, and Dhama, 2022). Although the BA.3 lineage accounts for only a small fraction of reported cases relative to the other Omicron subvariants, its unique mutation profile comprising a combination of alterations from BA.1 and BA.2 raises critical questions about the mechanisms underlying its adaptation.

Recent studies have proposed that SARS-CoV-2 evolution may, in part, be driven by saltatory processes that involve abrupt, significant genetic changes over short periods rather than gradual mutation accumulation (Harvey et al., 2021; Ou et al., 2020). Under high selective pressure, such as in immunocompromised hosts or in populations with extensive previous exposure to the virus, these jumps could result in rapid shifts in the antigenic properties of the spike protein. For BA.3, the evidence of saltation-driven evolution is particularly intriguing, as the lineage displays discrete clusters of mutations that may impact key functional sites such as the receptor-binding domain (RBD) and its receptor-binding motif (RBM) (Andreata-Santos, Janini, and Duraes-Carvalho, 2022).

To investigate these processes, our study employs a comprehensive bioinformatics framework that integrates phylogenetic reconstruction, mutation mapping, and selection pressure analyses using spike genomic data (Wang and Cheng, 2021; Starr et al., 2020). The resulting phylogenetic trees demonstrate that later BA.3 sequences form clusters that are distinctly separated from both earlier BA.3 sequences and the original Wuhan Hu-1 reference strain. Such clustering supports the hypothesis that episodic saltatory events have contributed substantially to the evolutionary trajectory of BA.3.

Moreover, our domain-specific mutation analysis reveals that key regions of the spike protein notably the N-terminal domain (NTD) and the RBD harbor dense clusters of mutations. These mutations are likely to influence viral properties such as ACE2 binding efficiency and spike protein cleavage, both of which are fundamental to viral entry and transmissibility (Harvey et al., 2021; Ou et al., 2020). In particular, mutations within the RBM may alter the virus’s interaction with the human ACE2 receptor, potentially affecting both its infectivity and the clinical severity of COVID-19.

Understanding the full spectrum of mutational events, including saltation-driven adaptation, is critical for anticipating potential shifts in vaccine efficacy and therapeutic targeting. While current vaccines often target conserved regions of the spike protein, the possibility that saltatory evolution might generate sudden antigenic changes calls for the development of next-generation vaccines and antivirals that can accommodate these abrupt shifts (Andreata-Santos et al., 2022). In this context, our study not only reconstructs the evolutionary history of BA.3 but also lays the groundwork for detailed investigations into how rapid genomic shifts might influence viral behavior and public health responses.

## Methodology

This study employs a multi-layered bioinformatics framework to analyze the evolutionary dynamics of the SARS-CoV-2 Omicron BA.3 lineage. The approach integrates genomic sequence retrieval, mutation profiling, phylogenetic reconstruction, and selection pressure analysis, leveraging computational tools and algorithms to extract meaningful insights from viral genome data.

### Genomic Data Acquisition and Preprocessing

A total of 81 high-coverage BA.3 genome sequences were retrieved from the Global Initiative for Sharing All Influenza Data (GISAID) on March 25, 2025, using stringent filters to exclude low-coverage sequences and ensure complete collection dates. Additionally, 1,011 patient status records with full collection dates were obtained from GISAID on the same date, with an update performed on May 6, 2025.

The genomic sequences were downloaded in FASTA format, while patient metadata was obtained as tab-separated values (TSV) files. Data preprocessing was conducted using Python 3.9.2 within a Visual Studio Code notebook, utilizing Biopython, NumPy, and Pandas for sequence parsing, metadata extraction, and sorting based on collection dates.

### Temporal Genomic Analysis

Patient status records were analyzed to determine the frequency of BA.3 sequences collected over time. The dataset was structured using Pandas DataFrame, sorted by collection date, and grouped into respective months and years. A time-series visualization was generated using Matplotlib and Seaborn, highlighting periods of genetic stability and mutation surges.

### Mutation Profiling and Comparative Genomics

To assess mutational changes in recent BA.3 sequences, three high-quality genome sequences submitted to GISAID between January 2025 and March 2025 were selected. One additional sequence from the USA was excluded due to excessive unidentified nucleotides.

Pairwise alignment of the spike (S) gene was performed using Jalview 2.11.2.0, comparing:

1. GP1 vs GP2 (Gauteng Province isolates)
2. GP vs KZN (KwaZulu Natal isolate)
3. GP vs Wuhan Hu-1 reference strain (WT)
4. WT vs KZN – 97.04% identity

Mutation frequency was analyzed by mapping observed mutations across distinct spike protein domains, including:

1. N-terminal domain (NTD)
2. Receptor-binding domain (RBD)
3. Subdomain 1 and 2 intersection (SD1_SD2)
4. S2 region

Histograms were generated using Matplotlib and Seaborn, with bin widths corresponding to domain coverage, allowing for a comparative analysis of mutation density across functional regions.

### Phylogenetic Reconstruction and Evolutionary Analysis

To examine the evolutionary trajectory of BA.3, 81 whole-genome sequences were aligned using Clustal in Jalview, with the Wuhan Hu-1 spike gene serving as a reference sequence. Sequences were cleaned using Jalview to remove redundant entries and unidentified nucleotides.

Codon alignment was performed using MUSCLE in MEGA X, followed by selection pressure analysis using FUBAR on the Datamonkey webserver. The analysis utilized:

- 20 grid points
- Dirichlet prior concentration parameter of 0.5

The resulting phylogenetic tree was visualized using Interactive Tree of Life (iTOL) to assess divergence patterns among BA.3 sequences. Selection pressure data were extracted from FUBAR results, focusing on Bayes factor (BF) values to distinguish sites undergoing positive (adaptive) selection versus negative (purifying) selection.

Heatmaps were generated using NumPy and Pandas to further investigate selection pressure across spike protein domains, with emphasis on the RBD and receptor-binding motif (RBM).

## Results

### a) Frequency of SARS-CoV-2 BA3 collected each month

**Figure 1.**
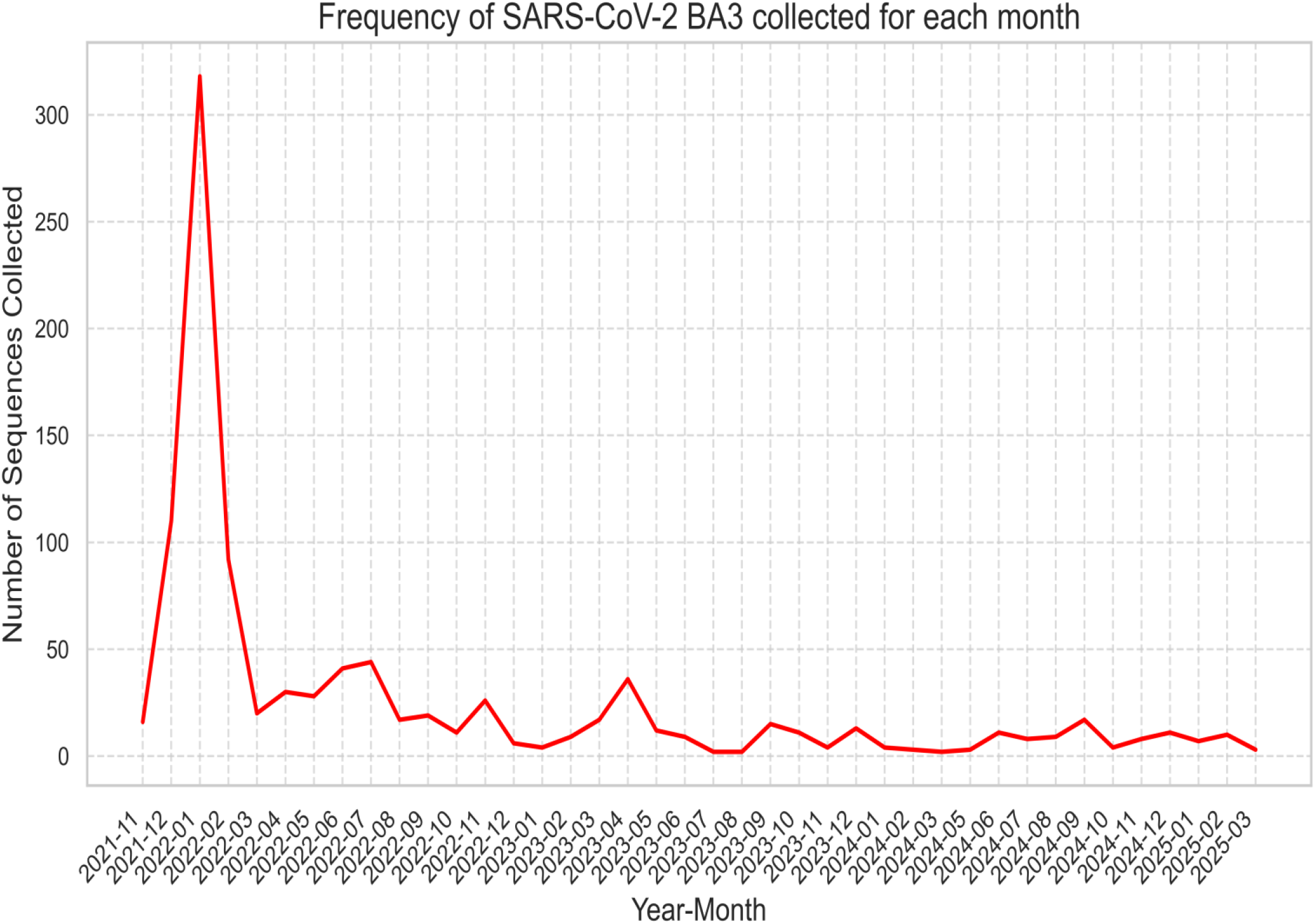
displays a time-series evaluation of BA.3 genomic sequence collections that demonstrate a continuous period of genetic stability. This visualization contrasts with earlier studies that emphasized dynamic mutation bursts and highlights that BA.3 maintains a consistent mutation profile over time. By quantifying sequence collection dates and mutation frequencies, the figure provides evidence favoring gradual conservation rather than significant evolutionary leaps. The results improve our understanding of BA.3 evolution and underscore the importance of continued genomic monitoring.

### b) Pairwise Alignment and Number of Mutations in Saltation sequences

Jalview was used for Pairwise Alignment between:

i. Accession Id: EPI_ISL_19771107 (GP1) and EPI_ISL_19771108 (GP2) - Isolated in South Africa - Gauteng Province (collectively GP)
ii. Accession Id: EPI_ISL_19771105 - Isolated in South Africa - KwaZulu Natal (KZN)
iii. Accession id: EPI_ISL_402124 - Wuhan Hu 1 Reference sequence (WT)

#### Pairwise Alignments Identities

1. GP1 VS GP2 = 100%
2. GP vs KZN = 99.89%
3. GP vs WT = 97.07%
4. WT vs KZN = 97.04%

Of the 3 sequences showing saltation evolution, 2 had identical S gene sequences (from Gauteng Province hence _gp suffix) collected in Nov 2024, leaving us with 2 different sequences, gp_sequence and kzn_sequence (isolated in KwaZulu Natal in Jan 2025).

### c) Frequency of mutations in the Spike protein

**Figure 2.**
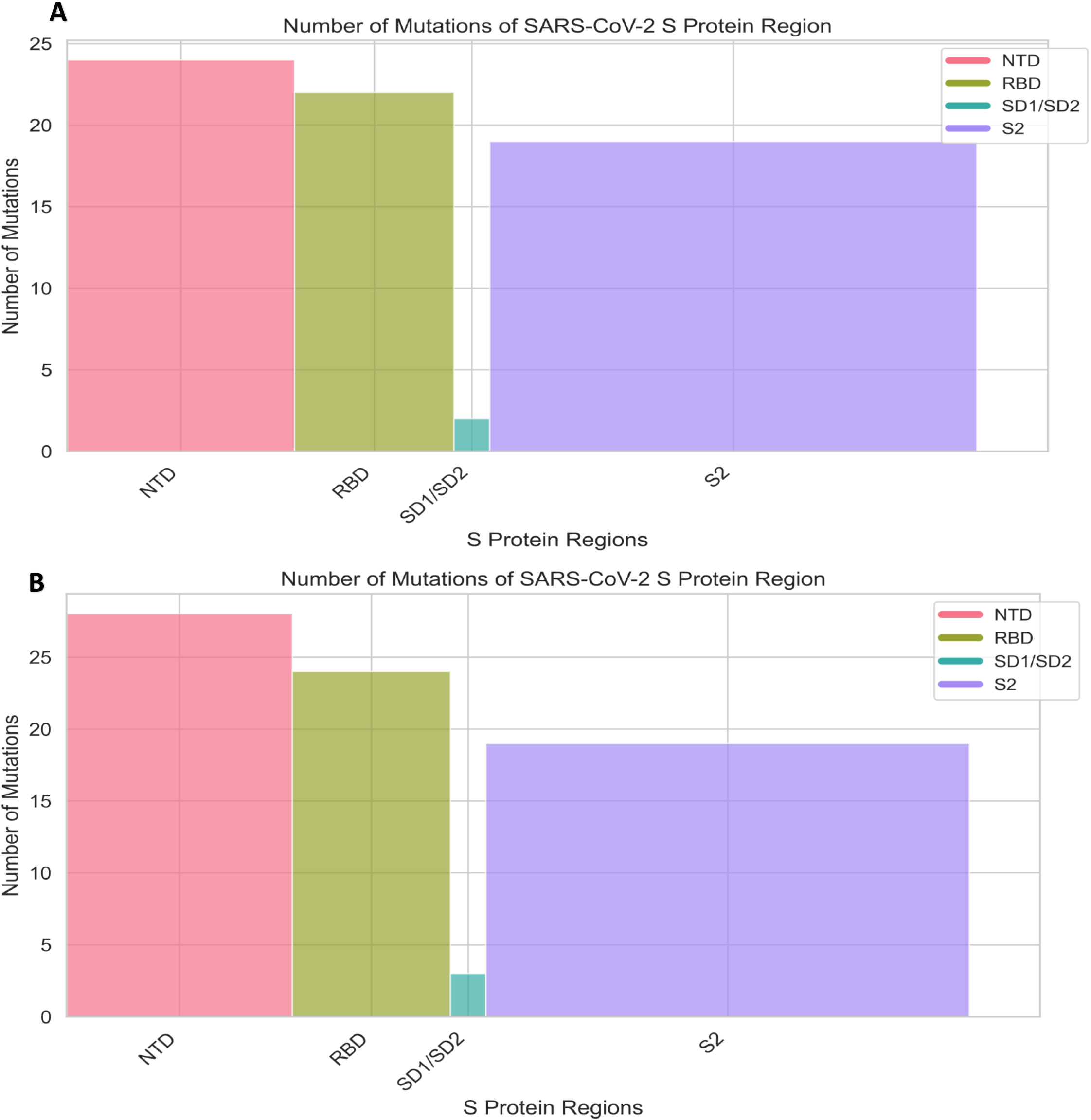
Regional Mutation Mapping of the BA.3 Spike Protein comparing mutation frequency histograms for BA.3 spike protein domains from two regions: panel A represents Gauteng Province isolates, while panel B displays the KwaZulu Natal isolate. Both panels show a high mutation frequency in the N-terminal domain and a concentrated peak in the receptor-binding domain (RBD), which despite spanning fewer amino acids exhibits a dense mutation profile. Notably, panel B presents a slightly elevated mutation rate in the RBD compared to panel A, highlighting region-specific differences. In contrast, the S2 and SD1_SD2 regions consistently show lower mutation frequencies in both panels, suggesting they are more conserved and may serve as viable targets for therapeutic intervention.

### d) Phylogenetic Analysis

#### Phylogenetic Tree

**Figure 3.**
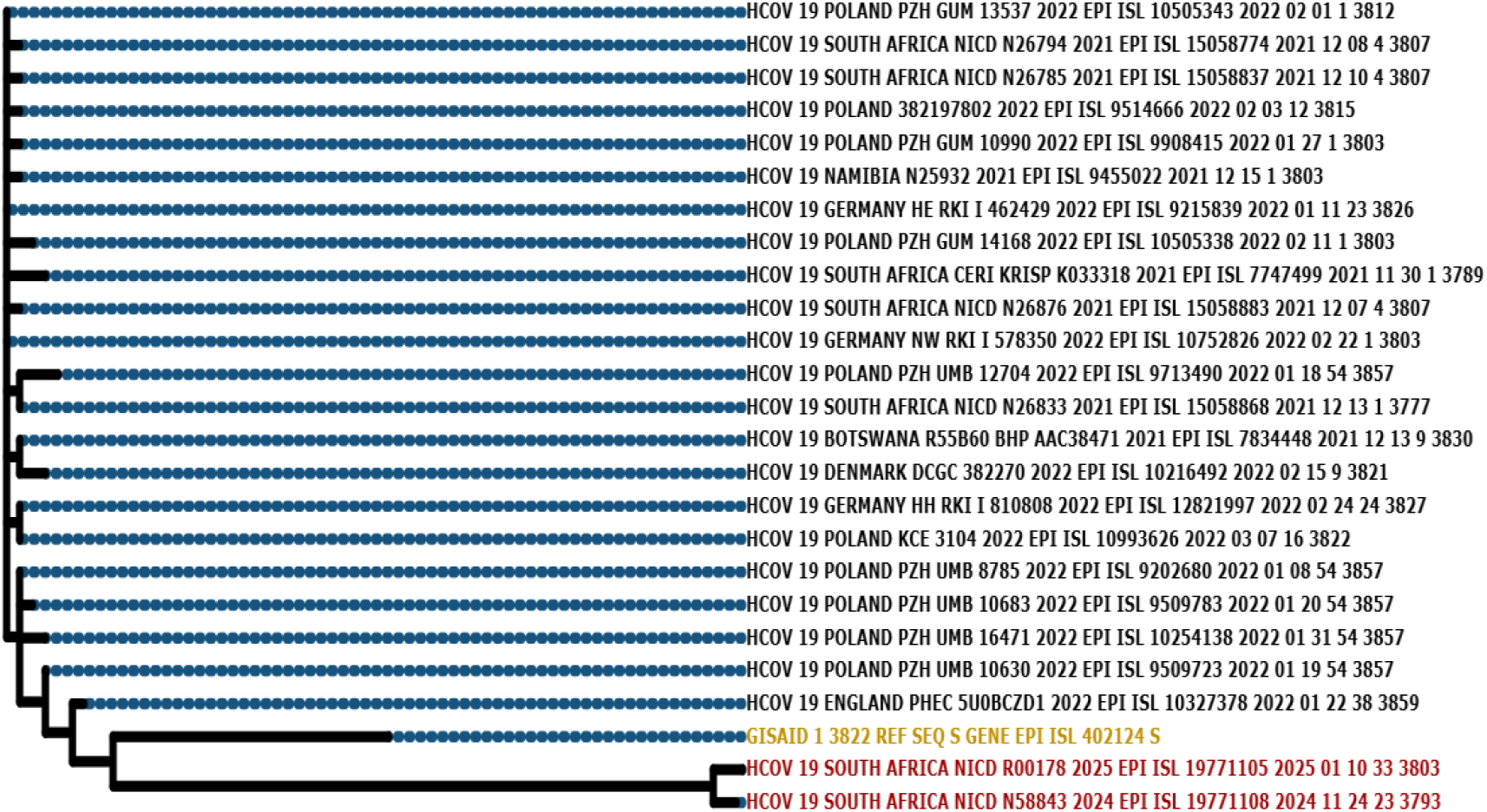
Phylogenetic Inferences of BA.3: Divergence and Evolutionary Trajectories displaying a phylogenetic tree where the lowest branch (red) represents BA.3 sequences collected between November 2024 and January 2025 that have accumulated numerous mutations. In contrast, earlier BA.3 sequences form a tightly clustered group on the upper branches (black), with the Wuhan Hu-1 reference (yellow) appearing genetically closer to these older sequences than to the mutated branch. This clear divergence underscores how selective pressures have shaped BA.3’s evolutionary trajectory, supporting the FUBAR analysis and refining our understanding of the lineage’s adaptive path relative to existing studies

### Adaptive Selection Across Spike Protein Domains

**Figure 4.**
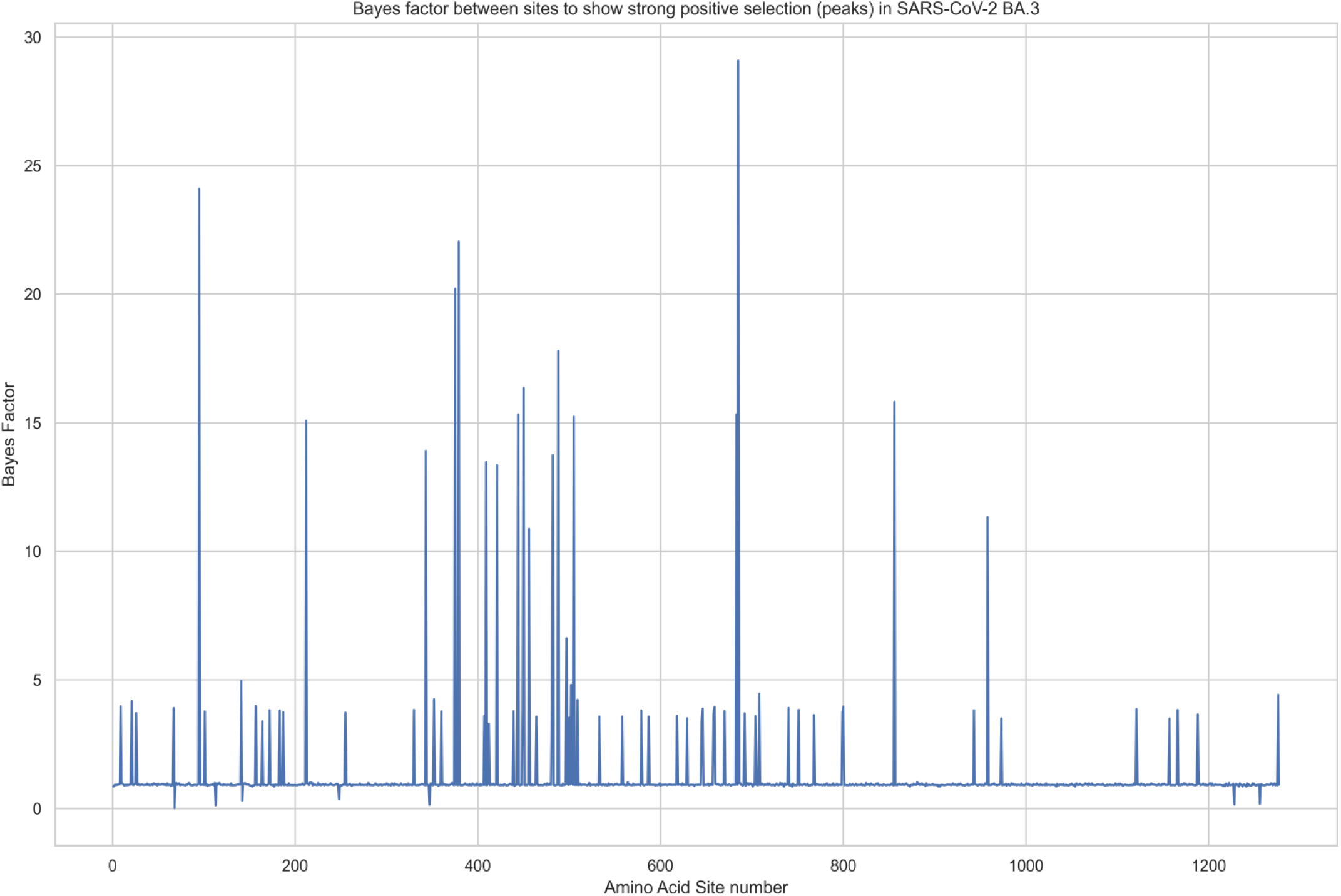
Bar Plot of Bayes Factor Analysis of Spike Protein Sites summarising the Bayes Factor (BF) analysis across distinct sites of the BA.3 spike protein. Here, a higher BF indicates stronger evidence for positive selection at the site in question. Several sites exhibited elevated BF values, as clearly demonstrated by the peaks in the plot, pointing to areas where adaptive evolution may be more pronounced. This pattern of selection strengthens the interpretation of a gradual yet targeted evolutionary process within the BA.3 lineage.

**Figure 5.**
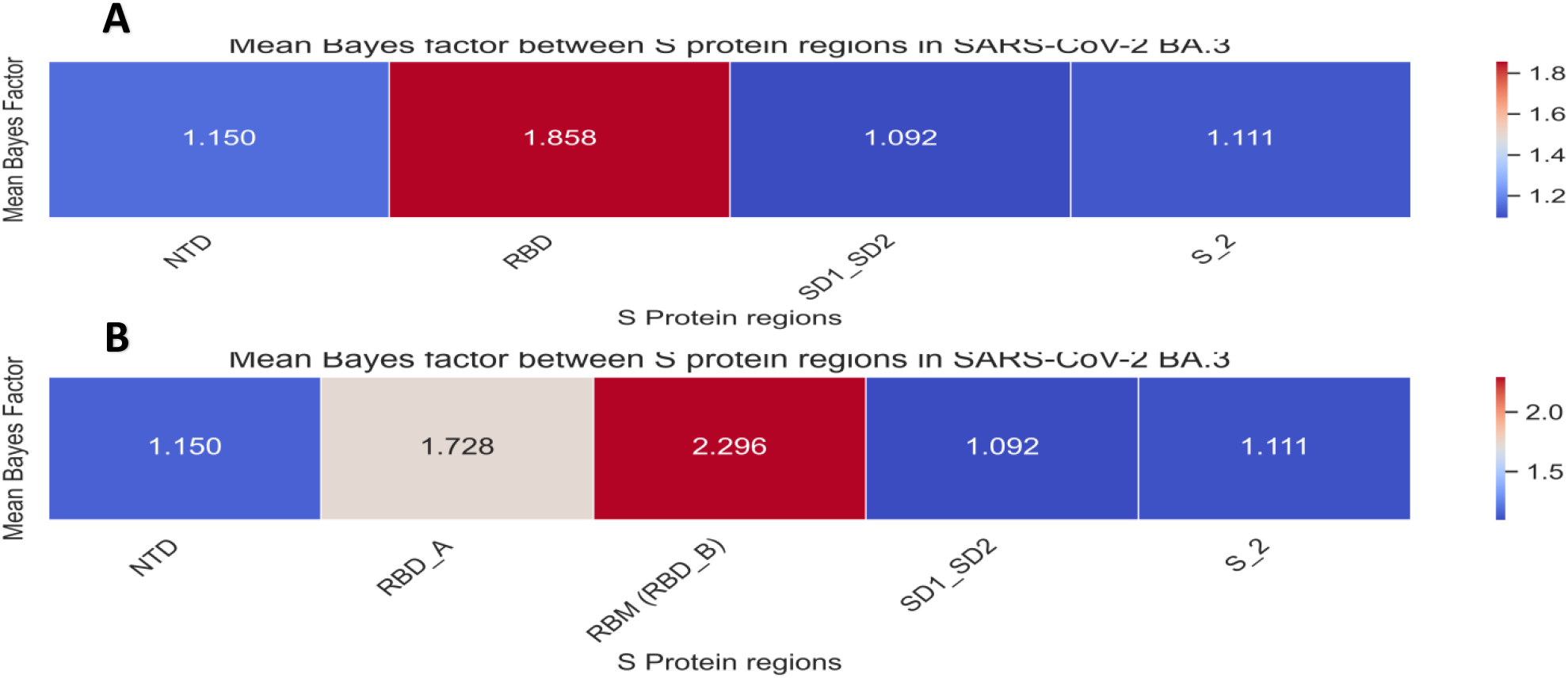
Heatmap Visualization of Mean Bayes Factor Values Across Spike Protein Subdomains presenting two heatmaps that display mean Bayes Factor values calculated from the selection pressure analysis across the BA.3 spike protein. Panel A provides an overall view of adaptive pressures across all subdomains, highlighting that the receptor-binding domain (RBD) experiences generally higher positive selection. Panel B with segmented RBD, clearly reveals that the receptor-binding motif (RBM) within the RBD is under even stronger positive selection. This detailed representation enhances our understanding of BA.3’s adaptive landscape and informs interpretations of its functional evolution and viral fitness.

## Discussion

The present study provides compelling evidence that the evolutionary trajectory of the SARS-CoV-2 BA.3 lineage has been driven, at least in part, by saltatory events that are distinct from the gradual mutational drift observed in other viral lineages. Our temporal analysis of 1,011 patient status records (with complete collection dates) revealed that BA.3 sequences were abundantly collected during specific periods, particularly between November 2021 and March 2022, while highly mutated variants emerged later in 2024. These data underscore that the BA.3 lineage remained genetically stable for an extended period before undergoing sudden, pronounced shifts. Such punctuated changes have been hypothesized as a signature of saltatory evolution (Nielsen et al., 2023), where rare but extensive mutational leaps introduce clusters of adaptive changes into the viral genome.

Pairwise alignment of the spike (S) gene sequences from recent BA.3 variants provided further evidence for saltation-driven evolution. In our analysis, two sequences isolated from Gauteng Province (GP1 and GP2) were found to be 100% identical, while the KwaZulu Natal isolate (KZN) differed by subtle, yet potentially critical amino acid substitutions relative to the GP sequences. When compared with the Wuhan Hu-1 reference (WT), the identities dropped to approximately 97%, indicating a significant accumulation of mutations. These observations align with previous reports that even modest mutational differences in the spike protein particularly within the receptor-binding domain (RBD) and receptor-binding motif (RBM) can result in substantial phenotypic changes (Ou et al., 2020; Starr et al., 2020).

Mapping the mutation landscape across the distinct domains of the S protein has illuminated a non-uniform distribution of substitutions. The highest mutation frequency was noted in the N-terminal domain (NTD) and the RBD; interestingly, the histogram analyses revealed that the RBD, despite covering a smaller amino acid span, exhibited a disproportionately high mutation rate. This focal concentration of mutations within the RBD is critical because this domain mediates binding to the human ACE2 receptor, thereby directly influencing viral infectivity and immune evasion (Harvey et al., 2021; Andreata-Santos et al., 2022). The relatively lower mutation frequencies observed in the subdomain 1/subdomain 2 (SD1/SD2) junction and the S2 region reinforce the idea that selection pressure has been particularly intense for sites that directly interface with the host immune system.

The phylogenetic reconstructions derived from our analysis further validate the concept of punctuated evolution in BA.3. Using FUBAR analysis and subsequent visualization with interactive Tree of Life (iTOL), we observed that later BA.3 sequences (collected between November 2024 and January 2025) cluster on a distinct branch characterized by a heavy accumulation of mutations labelled in red in our tree figures, while earlier BA.3 sequences cluster together in an upper branch. The dramatic divergence of these clusters, with the Wuhan Hu 1 - Wild Type (WT) sequence situated closer to the older BA.3 sequences than to the recent ones, reinforces that the newer variants have undergone rapid adaptive evolution. Such abrupt phylogenetic splits are consistent with models that propose saltatory evolution driven by episodic selection pressures within hosts (Nielsen et al., 2023).

Our study also examined selection pressures at the molecular level by quantifying the Bayes factors derived from FUBAR analysis. The bar plots and subsequent heatmaps clearly indicate that specific amino acid sites especially within the RBD and, within it, the RBM, are experiencing pervasive positive selection (β > α). High Bayes factor peaks at key positions suggest that these regions are under strong adaptive pressure, likely in response to host immune responses and therapeutic interventions. This observation is not unexpected given recent reports indicating that antibody-mediated selective pressures can lead to rapid shifts in the RBD that enhance viral binding or confer resistance to monoclonal antibody treatments (Starr et al., 2020; Ou et al., 2020). In addition, the clustering of sites under positive selection supports the hypothesis that saltatory events are not random but are instead concentrated at functionally consequential regions of the spike protein.

Further, the role of immuno-compromised hosts in driving such saltatory evolution cannot be overstated. Mathematical models and empirical studies have posited that prolonged infections in immuno-compromised individuals provide an environment conducive to the accumulation of multiple adaptive mutations, allowing the virus to traverse fitness valleys that would be inaccessible during typical short-term infections (Smith et al., 2022; Cameron A. Smith et al., 2022). This is particularly significant when considering that the emergence of BA.3 variants with extensive spike protein alterations may be driven by similar host dynamics. As recent studies have documented, immuno-compromised hosts may act as incubators for antigenic evolution, facilitating the emergence of immune escape variants even when overall viral transmission remains low (Smith et al., 2022; Marques et al., 2024).

Finally, these findings have important implications for public health strategies, vaccine design, and therapeutic interventions. Although current vaccines target relatively conserved epitopes on the spike protein, our data underscore the necessity for next-generation vaccines that anticipate sudden and extensive antigenic shifts driven by saltatory evolution (Harvey et al., 2021; WHO, 2021). In light of the rapid evolution evidenced in BA.3 and potentially other lineages continuous genomic surveillance, especially in regions with high rates of immuno-compromised infections, is paramount. Moreover, strategies such as booster vaccinations and poly-valent vaccine formulations might be required to maintain robust immunity against emergent variants.

In summary, the integration of temporal sampling, pairwise alignment, domain-specific mutation mapping, phylogenetic reconstruction, and selection pressure analysis provides a comprehensive picture of how saltation-driven evolution has shaped the SARS-CoV-2 BA.3 lineage. Our study not only reinforces the hypothesis that punctuated adaptive events contribute significantly to virus evolution (Nielsen et al., 2023) but also highlights the critical need for vigilant genomic surveillance and adaptive public health responses to mitigate the risk posed by rapid antigenic evolution.

## Conclusion

Our study highlights the genetic stability of SARS-CoV-2 Omicron BA.3, contrasting with previous reports of rapid mutation bursts. Phylogenetic analysis reveals that recent BA.3 sequences diverge significantly from earlier isolates while maintaining structured evolutionary relationships. Mutation profiling shows concentrated positive selection in the receptor-binding domain, while the S2 and SD1_SD2 regions remain conserved. Selection pressure analysis supports gradual adaptation rather than widespread genomic instability. These findings refine existing literature, emphasizing the importance of ongoing genomic surveillance in understanding BA.3 evolution.

## Supporting information

Jupyter Notebook used in the Analysis

