## Supplementary material for "Integrative genomic study of mutation dynamics and Evolutionary trends in SARS-CoV-2 omicron BA.3": Jupyter Notebook used in the Analysis

The Evolutionary trajectory of SARS-CoV-2 lineage BA.3\_Evidence for saltation-driven Adaptation


### The Evolutionary trajectory of SARS-CoV-2 lineage BA.3: Evidence for saltation-driven Adaptation¶

#### 1. Introduction¶

The SARS-CoV-2 pandemic has been characterised by the rapid emergence and spread of numerous variants. Among these, the SARS-CoV-2 BA.3 lineage of the omicron variant exhibited a distinct evolutionary trajectory, raising questions about the mechanisms driving its adaptation. This research project aims at investigating the evolutionary history of SARS-CoV-2 BA.3, specifically focussing on the hypothesis of saltation-driven adaptations. Saltation evolution, suggests rapid and substantial evolutionary changes occuring over short periods, potentially explaining the unique characteristics of SARS-CoV-2 BA.3.

#### 2. Research Question¶

Do saltation events play a significant role in the evolutionary adaptation of the SARS-CoV-2 BA.3 lineage?

#### 3. Objectives¶

> - To reconstruct the phylogenetic histroy of the SARS-CoV-2 BA.3 lineage using Spike genomic data. ✔
> - To identify and characterize the specific genomic mutations associated with the emergence and diversification of SARS-CoV-2 BA.3 new sequences.✔
>
> - To analyse the selective pressures acting on the SARS-CoV-2 BA.3 lineage, with a focus on detecting evidence of positive selection. ✔
> - To provide evidence for the saltation driven adaptations of the SARS-CoV-2 Ba.3 lineage. ✔
> - To investigate the potential structural and functional consequences of key mutations, particularly in the spike protein.
> - Optimise and design plots to match publication standards.
> - Discussing results in detail.

#### 4. Methodology¶

This project employs a comprehensive bioinformatics approach, integrating phylogenetics, genomic data science, structural and functional proteomics. Key methdologies include:

> - **Phylogenetic Analysis** : Construction of Nucleotide GTR (General Time Reversible) model hosted on Datamonkey - FUBAR webserver.
> - **Genomic Variation Analysis** : Divided into 2 parts
>
> 1. Identification of unique mutations using Python in-built packaages and
> 2. dN/dS ratio calculations to assess selective pressures using FUBAR and Panel Dataframes (Pandas) Python data science package for calculations and loading result data. Matplotlib and Seaborn used for Data VIsualisation.

##### Sequence acquisition and extraction of the Spike (S) gene from whole genome sequences.¶

THe research followed 2 pathways:

1. Analysis of S genes from genomic sequences showing saltatory evolution against the Wuhan Hu 1 - S gene as reference sequence (n = 3).

   > - Patient status data was downloaded too, to determine mutations unique to the saltatory amino acid sequences using Wuhan Hu 1 S protein as reference
2. Analysis of all available BA.3 to calculate sites under selection pressures in all BA.3 sequences especially ones being driven towards positive selection. (n = 81), used a filter to only download sequences with `low coverage excluded`. Ended up with 64 sequences after cleaning and removing redundant sequences.

> - Resultant tree from FUBAR analysis was used for drawing the Phylogenetic tree to show how the novel BA.3 are distant from their cousins.

- *Anomaly detected during Multiple sequences alignment where 1 sequence (Accession ID:11120428) had a repeat on nucleotide (nt) position 2323-2336 was mirrored from nt position 2309-2322*

##### Sequence Analysis¶

In [5]:

```
# Python packages to be used in the project

import numpy as np

from scipy import stats

# For loading and displaying .csv files as DataFrames (Tables)
import pandas as pd

# Biopython is used to interact wuth biological data in Python Language
import Bio

from collections import Counter  #in-built python useful for counting items

# for data visualisation
import matplotlib.pyplot as plt
import seaborn as sns
```

In [6]:

```
from Bio import SeqIO       #importing the function for handling sequences from a file

ba3_spike = []                  # the variable holds the spike sequences to be excised

for seq_record in SeqIO.parse('Saltation ones.fasta','fasta'):         # loads sequences in 'anommalous file'
    spike_gene = seq_record[21500:25499]                               # excises approximate S gene genomic locus for each sequence and stored in spike gene variable
    ba3_spike.append(spike_gene)  
    print(seq_record.id)
                                         # each excised S gene is loaded into the ba3.spike dictionary defined above


# SeqIO.write(ba3_spike,'BA3 Saltation Spikes.fasta','fasta')          # method writes all spikes in ba3_spikes dictionary into a file named BA3 Saltation Spikes.fasta
#above statement commented to avoid overwriting the file wheneve code is executed.
```

```
hCoV-19/South_Africa/NICD-R00178/2025|EPI_ISL_19771105|2025-01-10
hCoV-19/South_Africa/NICD-N58822/2024|EPI_ISL_19771107|2024-11-22
hCoV-19/South_Africa/NICD-N58843/2024|EPI_ISL_19771108|2024-11-24
hCoV-19/USA/CO-CDPHE-43093479/2025|EPI_ISL_19775777|2025-02-26
```

THe USA sequence was discarded because it had many unidentified nucleotides which disrupts downstream processes.

In [7]:

```
from Bio import SeqIO                             #same method as above only that in this code block, S genes for all downloaded BA.3 whole 
                                                    #genome sequences are being excised.
ba3_spike = []

for seq_record in SeqIO.parse('BA3_low coverage excluded.fasta','fasta'):
    spike_gene=seq_record[21500:25499]
    ba3_spike.append(spike_gene)


# SeqIO.write(ba3_spike, 'BA3 All Spikes.fasta','fasta' )  
# Line was deactivated to avoid overwriting the files whenever the code is executed
```

##### Patient Status Analysis¶

Patient status data was downloaded from GISAID in .tsv (tab-seperated values), an online tool https://www.convertsimple.com was used to convert .tsv to .csv (comma-seperated values). *.csv* files are easy to handle and manipulate using existing Data Science packages than .tsv files.

THe Main reason for dowloading Patient Data Status was to investigate collection dates and amino acid changes.

In [8]:

```
import pandas as pd

ba3_patient_status = pd.read_table('ba3_May update patient status.tsv')
ba3_patient_status.tail()
```

Out[8]:

|  | Virus name | Accession ID | Collection date | Location | Host | Additional location information | Sampling strategy | Gender | Patient age | Patient status | Last vaccinated | Passage | Specimen | Additional host information | Lineage | Clade | AA Substitutions |
| --- | --- | --- | --- | --- | --- | --- | --- | --- | --- | --- | --- | --- | --- | --- | --- | --- | --- |
| 1007 | hCoV-19/Poland/PZH-UMB-13810/2022 | EPI\_ISL\_9977822 | 2022-02-01 | Europe / Poland / Podlaskie / Lomza | Human | NaN | NaN | Male | 73 | unknown | NaN | Original | NaN | NaN | BA.3 | GRA | (NSP5\_P132H,Spike\_H69del,Spike\_T95I,NSP6\_A88V,... |
| 1008 | hCoV-19/Poland/PZH-UMB-14004/2022 | EPI\_ISL\_9978008 | 2022-02-03 | Europe / Poland / Podlaskie / Sokolski | Human | NaN | NaN | Male | 37 | unknown | NaN | Original | NaN | NaN | BA.3 | GRA | (NSP5\_P132H,Spike\_H69del,Spike\_T95I,NSP6\_A88V,... |
| 1009 | hCoV-19/Poland/PZH-UMB-14211/2022 | EPI\_ISL\_9978209 | 2022-01-26 | Europe / Poland / Malopolskie / Krakow | Human | NaN | NaN | Female | 31 | unknown | NaN | Original | NaN | NaN | BA.3 | GRA | (NSP5\_P132H,Spike\_H69del,Spike\_T95I,NSP6\_A88V,... |
| 1010 | hCoV-19/England/MILK-367CA02/2022 | EPI\_ISL\_9989592 | 2022-02-11 | Europe / United Kingdom / England | Human | NaN | NaN | unknown | unknown | unknown | NaN | Original | NaN | NaN | BA.3 | GRA | (NSP5\_P132H,Spike\_H69del,Spike\_T95I,NSP6\_A88V,... |
| 1011 | hCoV-19/Scotland/QEUH-36491AF/2022 | EPI\_ISL\_9996088 | 2022-02-10 | Europe / United Kingdom / Scotland | Human | NaN | NaN | unknown | unknown | unknown | NaN | Original | NaN | NaN | BA.3 | GRA | (NSP5\_P132H,Spike\_H69del,Spike\_T95I,NSP6\_A88V,... |

##### Collection Date Analysis¶

TO understand how the number of sequences which were collected during the course of BA.3, a line graph was plotted of number of collected BA.3 viruses against dates of collection.

In [9]:

```
from datetime import datetime, date
collection_dates_series = ba3_patient_status['Collection date']

collection_dates_list = list(collection_dates_series)      # taking collection dates from panda

# collection_dates = [datetime.strptime(date, '%Y-%m-%d').date() for date in collection_dates_list] 

collection_dates = [datetime.strptime(item, '%Y-%m-%d').date() if isinstance(item,str) else item for item in collection_dates_list] 
# converting all adtes to date format
print(collection_dates)
```

```
[datetime.date(2022, 2, 3), datetime.date(2022, 1, 29), datetime.date(2022, 2, 8), datetime.date(2022, 2, 3), datetime.date(2022, 2, 7), datetime.date(2022, 2, 6), datetime.date(2022, 2, 13), datetime.date(2022, 2, 12), datetime.date(2022, 2, 2), datetime.date(2022, 1, 27), datetime.date(2022, 1, 28), datetime.date(2022, 1, 29), datetime.date(2022, 1, 28), datetime.date(2022, 1, 31), datetime.date(2022, 1, 31), datetime.date(2022, 1, 31), datetime.date(2022, 2, 4), datetime.date(2022, 2, 6), datetime.date(2022, 2, 2), datetime.date(2021, 12, 4), datetime.date(2021, 12, 8), datetime.date(2021, 12, 8), datetime.date(2022, 1, 2), datetime.date(2022, 1, 7), datetime.date(2022, 1, 11), datetime.date(2022, 1, 8), datetime.date(2022, 2, 15), datetime.date(2022, 1, 31), datetime.date(2022, 2, 1), datetime.date(2022, 2, 2), datetime.date(2022, 2, 4), datetime.date(2022, 2, 4), datetime.date(2022, 2, 3), datetime.date(2022, 2, 3), datetime.date(2022, 2, 4), datetime.date(2022, 2, 3), datetime.date(2022, 2, 3), datetime.date(2022, 1, 31), datetime.date(2022, 1, 27), datetime.date(2022, 2, 1), datetime.date(2022, 2, 1), datetime.date(2022, 2, 7), datetime.date(2022, 2, 8), datetime.date(2022, 2, 10), datetime.date(2022, 2, 12), datetime.date(2022, 2, 14), datetime.date(2022, 1, 22), datetime.date(2022, 2, 21), datetime.date(2022, 2, 9), datetime.date(2022, 2, 9), datetime.date(2022, 2, 13), datetime.date(2022, 2, 18), datetime.date(2022, 2, 7), datetime.date(2022, 2, 6), datetime.date(2021, 12, 6), datetime.date(2021, 12, 24), datetime.date(2022, 2, 14), datetime.date(2022, 2, 14), datetime.date(2022, 2, 11), datetime.date(2022, 2, 1), datetime.date(2022, 2, 5), datetime.date(2022, 2, 28), datetime.date(2022, 2, 15), datetime.date(2022, 2, 1), datetime.date(2022, 2, 9), datetime.date(2022, 2, 3), datetime.date(2022, 1, 21), datetime.date(2021, 12, 1), datetime.date(2021, 12, 7), datetime.date(2022, 2, 14), datetime.date(2022, 2, 16), datetime.date(2022, 2, 28), datetime.date(2022, 2, 8), datetime.date(2022, 2, 15), datetime.date(2022, 2, 17), datetime.date(2022, 2, 18), datetime.date(2022, 2, 8), datetime.date(2022, 2, 28), datetime.date(2022, 2, 15), datetime.date(2022, 2, 16), datetime.date(2022, 2, 16), datetime.date(2022, 2, 13), datetime.date(2022, 2, 22), datetime.date(2022, 1, 22), datetime.date(2022, 2, 17), datetime.date(2022, 2, 20), datetime.date(2022, 2, 17), datetime.date(2022, 2, 17), datetime.date(2022, 1, 5), datetime.date(2021, 12, 9), datetime.date(2021, 12, 9), datetime.date(2021, 12, 6), datetime.date(2022, 2, 4), datetime.date(2022, 3, 7), datetime.date(2022, 2, 9), datetime.date(2022, 3, 7), datetime.date(2022, 3, 7), datetime.date(2022, 1, 18), datetime.date(2021, 12, 28), datetime.date(2021, 12, 31), datetime.date(2021, 12, 31), datetime.date(2021, 12, 31), datetime.date(2021, 12, 31), datetime.date(2021, 12, 20), datetime.date(2021, 12, 20), datetime.date(2021, 12, 20), datetime.date(2022, 1, 28), datetime.date(2022, 2, 21), datetime.date(2022, 2, 22), datetime.date(2022, 2, 22), datetime.date(2022, 2, 22), datetime.date(2021, 12, 1), datetime.date(2022, 1, 28), datetime.date(2022, 1, 28), datetime.date(2022, 2, 21), datetime.date(2022, 3, 15), datetime.date(2022, 2, 24), datetime.date(2021, 12, 25), datetime.date(2021, 12, 6), datetime.date(2022, 2, 21), datetime.date(2022, 1, 20), datetime.date(2022, 3, 4), datetime.date(2022, 2, 20), datetime.date(2022, 2, 18), datetime.date(2022, 2, 20), datetime.date(2022, 2, 4), datetime.date(2022, 3, 16), datetime.date(2022, 3, 15), datetime.date(2022, 2, 21), datetime.date(2022, 3, 22), datetime.date(2022, 3, 23), datetime.date(2022, 3, 23), datetime.date(2022, 2, 14), datetime.date(2022, 3, 23), datetime.date(2022, 3, 25), datetime.date(2022, 3, 26), datetime.date(2022, 3, 10), datetime.date(2022, 1, 5), datetime.date(2021, 12, 24), datetime.date(2021, 12, 2), datetime.date(2021, 12, 6), datetime.date(2021, 12, 9), datetime.date(2021, 12, 9), datetime.date(2021, 12, 9), datetime.date(2021, 12, 10), datetime.date(2021, 12, 10), datetime.date(2021, 12, 10), datetime.date(2021, 12, 6), datetime.date(2021, 12, 6), datetime.date(2021, 12, 10), datetime.date(2021, 12, 10), datetime.date(2022, 4, 1), datetime.date(2022, 3, 16), datetime.date(2022, 3, 23), datetime.date(2022, 3, 14), datetime.date(2022, 1, 27), datetime.date(2022, 1, 28), datetime.date(2022, 1, 30), datetime.date(2022, 1, 30), datetime.date(2022, 1, 31), datetime.date(2022, 2, 1), datetime.date(2022, 1, 23), datetime.date(2022, 1, 30), datetime.date(2022, 1, 19), datetime.date(2022, 1, 25), datetime.date(2022, 4, 17), datetime.date(2022, 4, 16), datetime.date(2022, 4, 21), datetime.date(2022, 4, 20), datetime.date(2022, 4, 15), datetime.date(2022, 4, 16), datetime.date(2022, 4, 7), datetime.date(2022, 4, 20), datetime.date(2022, 1, 27), datetime.date(2022, 1, 30), datetime.date(2022, 1, 29), datetime.date(2022, 1, 30), datetime.date(2022, 4, 23), datetime.date(2022, 2, 15), datetime.date(2022, 2, 7), datetime.date(2022, 1, 31), datetime.date(2022, 4, 12), datetime.date(2022, 3, 11), datetime.date(2022, 5, 1), datetime.date(2022, 4, 28), datetime.date(2022, 2, 24), datetime.date(2022, 2, 14), datetime.date(2022, 5, 6), datetime.date(2022, 4, 25), datetime.date(2022, 3, 16), datetime.date(2022, 4, 25), datetime.date(2022, 4, 30), datetime.date(2022, 4, 19), datetime.date(2022, 4, 11), datetime.date(2022, 4, 7), datetime.date(2022, 4, 19), datetime.date(2022, 5, 17), datetime.date(2022, 5, 24), datetime.date(2022, 5, 27), datetime.date(2022, 5, 28), datetime.date(2022, 5, 11), datetime.date(2022, 2, 4), datetime.date(2022, 5, 11), datetime.date(2022, 6, 7), datetime.date(2022, 6, 6), datetime.date(2022, 6, 6), datetime.date(2022, 6, 8), datetime.date(2022, 6, 16), datetime.date(2022, 6, 16), datetime.date(2022, 6, 17), datetime.date(2022, 6, 19), datetime.date(2022, 5, 31), datetime.date(2022, 6, 14), datetime.date(2022, 6, 16), datetime.date(2022, 6, 16), datetime.date(2022, 6, 16), datetime.date(2022, 6, 16), datetime.date(2022, 6, 16), datetime.date(2022, 6, 16), datetime.date(2022, 5, 19), datetime.date(2022, 6, 13), datetime.date(2022, 6, 20), datetime.date(2022, 5, 2), datetime.date(2022, 6, 29), datetime.date(2022, 6, 12), datetime.date(2022, 6, 16), datetime.date(2022, 6, 25), datetime.date(2022, 6, 14), datetime.date(2022, 7, 1), datetime.date(2022, 7, 4), datetime.date(2022, 7, 8), datetime.date(2022, 6, 26), datetime.date(2022, 7, 2), datetime.date(2022, 7, 4), datetime.date(2022, 7, 8), datetime.date(2022, 7, 4), datetime.date(2022, 7, 4), datetime.date(2022, 7, 5), datetime.date(2022, 7, 5), datetime.date(2022, 7, 12), datetime.date(2022, 7, 13), datetime.date(2022, 7, 13), datetime.date(2022, 7, 2), datetime.date(2022, 7, 6), datetime.date(2022, 7, 7), datetime.date(2022, 7, 8), datetime.date(2022, 7, 10), datetime.date(2022, 7, 10), datetime.date(2022, 7, 6), datetime.date(2022, 6, 27), datetime.date(2022, 6, 17), datetime.date(2022, 6, 27), datetime.date(2022, 6, 26), datetime.date(2022, 6, 29), datetime.date(2022, 6, 7), datetime.date(2022, 7, 18), datetime.date(2022, 1, 19), datetime.date(2022, 7, 24), datetime.date(2022, 7, 24), datetime.date(2022, 7, 25), datetime.date(2022, 7, 21), datetime.date(2022, 7, 25), datetime.date(2022, 7, 18), datetime.date(2022, 7, 22), datetime.date(2022, 7, 25), datetime.date(2021, 12, 6), datetime.date(2021, 12, 6), datetime.date(2022, 7, 15), datetime.date(2022, 6, 14), datetime.date(2022, 4, 11), datetime.date(2021, 12, 6), datetime.date(2022, 8, 2), datetime.date(2022, 4, 6), datetime.date(2022, 4, 19), datetime.date(2022, 6, 23), datetime.date(2022, 1, 5), datetime.date(2022, 8, 13), datetime.date(2022, 8, 11), datetime.date(2022, 5, 24), datetime.date(2022, 6, 23), datetime.date(2021, 12, 7), datetime.date(2021, 12, 13), datetime.date(2021, 12, 21), datetime.date(2022, 6, 16), datetime.date(2021, 12, 8), datetime.date(2022, 8, 21), datetime.date(2022, 8, 23), datetime.date(2022, 8, 24), datetime.date(2022, 8, 23), datetime.date(2022, 8, 28), datetime.date(2022, 8, 26), datetime.date(2022, 8, 1), datetime.date(2022, 8, 2), datetime.date(2022, 8, 16), datetime.date(2022, 8, 16), datetime.date(2022, 9, 2), datetime.date(2021, 12, 8), datetime.date(2021, 12, 6), datetime.date(2021, 12, 10), datetime.date(2021, 12, 10), datetime.date(2021, 12, 13), datetime.date(2021, 12, 7), datetime.date(2021, 12, 15), datetime.date(2022, 9, 1), datetime.date(2022, 4, 23), datetime.date(2022, 4, 28), datetime.date(2022, 4, 29), datetime.date(2022, 5, 5), datetime.date(2022, 5, 7), datetime.date(2022, 5, 7), datetime.date(2022, 5, 10), datetime.date(2022, 6, 30), datetime.date(2022, 6, 22), datetime.date(2022, 9, 9), datetime.date(2022, 9, 10), datetime.date(2022, 9, 11), datetime.date(2022, 9, 4), datetime.date(2022, 9, 21), datetime.date(2022, 9, 12), datetime.date(2022, 9, 5), datetime.date(2022, 9, 8), datetime.date(2022, 9, 10), datetime.date(2022, 9, 12), datetime.date(2022, 9, 13), datetime.date(2022, 9, 13), datetime.date(2021, 12, 3), datetime.date(2022, 8, 7), datetime.date(2022, 7, 21), datetime.date(2022, 8, 31), datetime.date(2022, 5, 27), datetime.date(2022, 9, 10), datetime.date(2022, 9, 26), datetime.date(2022, 7, 25), datetime.date(2022, 10, 9), datetime.date(2022, 10, 11), datetime.date(2022, 10, 10), datetime.date(2022, 9, 17), datetime.date(2022, 7, 30), datetime.date(2022, 10, 18), datetime.date(2022, 10, 10), datetime.date(2022, 10, 12), datetime.date(2022, 10, 18), datetime.date(2022, 10, 19), datetime.date(2022, 10, 18), datetime.date(2022, 10, 18), datetime.date(2022, 11, 9), datetime.date(2022, 11, 9), datetime.date(2022, 10, 31), datetime.date(2022, 11, 5), datetime.date(2022, 11, 10), datetime.date(2022, 11, 12), datetime.date(2022, 11, 11), datetime.date(2022, 11, 1), datetime.date(2022, 11, 10), datetime.date(2022, 11, 22), datetime.date(2022, 5, 30), datetime.date(2022, 5, 25), datetime.date(2022, 7, 11), datetime.date(2022, 5, 24), datetime.date(2022, 4, 27), datetime.date(2022, 5, 8), datetime.date(2022, 5, 23), datetime.date(2022, 5, 20), datetime.date(2022, 5, 27), datetime.date(2022, 7, 20), datetime.date(2022, 7, 21), datetime.date(2022, 7, 20), datetime.date(2022, 8, 18), datetime.date(2022, 8, 15), datetime.date(2022, 9, 7), datetime.date(2022, 9, 3), datetime.date(2022, 5, 12), datetime.date(2022, 4, 29), datetime.date(2022, 5, 24), datetime.date(2022, 11, 13), datetime.date(2022, 11, 10), datetime.date(2022, 11, 15), datetime.date(2022, 11, 1), datetime.date(2022, 11, 2), datetime.date(2022, 5, 9), datetime.date(2022, 11, 1), datetime.date(2022, 11, 2), datetime.date(2022, 11, 4), datetime.date(2022, 11, 5), datetime.date(2022, 12, 7), datetime.date(2022, 11, 26), datetime.date(2022, 11, 25), datetime.date(2022, 4, 23), datetime.date(2022, 4, 13), datetime.date(2022, 12, 6), datetime.date(2023, 1, 2), datetime.date(2022, 12, 26), datetime.date(2023, 2, 6), datetime.date(2023, 2, 13), datetime.date(2023, 2, 12), datetime.date(2023, 2, 2), datetime.date(2023, 1, 1), datetime.date(2022, 12, 28), datetime.date(2022, 12, 4), datetime.date(2023, 3, 4), datetime.date(2023, 3, 3), datetime.date(2023, 2, 28), datetime.date(2023, 3, 31), datetime.date(2023, 3, 16), datetime.date(2023, 4, 2), datetime.date(2023, 3, 14), datetime.date(2023, 4, 1), datetime.date(2023, 3, 13), datetime.date(2023, 2, 22), datetime.date(2023, 4, 8), datetime.date(2023, 2, 27), datetime.date(2023, 3, 12), datetime.date(2023, 3, 12), datetime.date(2023, 3, 20), datetime.date(2022, 11, 29), datetime.date(2022, 11, 24), datetime.date(2023, 4, 19), datetime.date(2023, 1, 27), datetime.date(2023, 4, 4), datetime.date(2023, 3, 26), datetime.date(2023, 3, 11), datetime.date(2023, 3, 14), datetime.date(2023, 1, 23), datetime.date(2023, 4, 12), datetime.date(2023, 4, 13), datetime.date(2023, 4, 13), datetime.date(2023, 4, 17), datetime.date(2023, 4, 14), datetime.date(2022, 12, 3), datetime.date(2022, 11, 27), datetime.date(2022, 11, 28), datetime.date(2023, 4, 8), datetime.date(2023, 4, 13), datetime.date(2023, 4, 15), datetime.date(2023, 4, 19), datetime.date(2023, 4, 20), datetime.date(2023, 4, 21), datetime.date(2023, 4, 21), datetime.date(2023, 4, 24), datetime.date(2023, 4, 5), datetime.date(2023, 4, 15), datetime.date(2023, 3, 29), datetime.date(2023, 4, 11), datetime.date(2022, 11, 21), datetime.date(2022, 11, 4), datetime.date(2023, 5, 2), datetime.date(2023, 4, 11), datetime.date(2023, 4, 8), datetime.date(2023, 4, 16), datetime.date(2023, 4, 24), datetime.date(2023, 4, 21), datetime.date(2023, 5, 2), datetime.date(2023, 5, 4), datetime.date(2023, 3, 24), datetime.date(2023, 4, 13), datetime.date(2023, 4, 21), datetime.date(2023, 4, 25), datetime.date(2023, 5, 2), datetime.date(2023, 5, 4), datetime.date(2023, 5, 14), datetime.date(2023, 5, 17), datetime.date(2023, 5, 15), datetime.date(2023, 4, 15), datetime.date(2023, 5, 16), datetime.date(2023, 4, 15), datetime.date(2023, 4, 20), datetime.date(2023, 3, 20), datetime.date(2023, 5, 23), datetime.date(2023, 5, 1), datetime.date(2023, 6, 4), datetime.date(2023, 4, 26), datetime.date(2023, 4, 23), datetime.date(2023, 6, 1), datetime.date(2023, 6, 4), datetime.date(2023, 6, 10), datetime.date(2023, 6, 1), datetime.date(2023, 6, 9), datetime.date(2023, 6, 1), datetime.date(2023, 2, 3), datetime.date(2022, 5, 26), datetime.date(2023, 3, 29), datetime.date(2023, 7, 11), datetime.date(2023, 7, 27), datetime.date(2021, 12, 4), datetime.date(2023, 8, 29), datetime.date(2023, 9, 1), datetime.date(2023, 8, 21), datetime.date(2023, 9, 7), datetime.date(2023, 9, 11), datetime.date(2023, 9, 9), datetime.date(2023, 9, 15), datetime.date(2023, 9, 19), datetime.date(2023, 9, 21), datetime.date(2023, 9, 26), datetime.date(2023, 9, 26), datetime.date(2022, 7, 8), datetime.date(2022, 7, 9), datetime.date(2022, 7, 10), datetime.date(2023, 10, 4), datetime.date(2023, 10, 6), datetime.date(2023, 10, 3), datetime.date(2023, 9, 25), datetime.date(2022, 7, 29), datetime.date(2022, 7, 30), datetime.date(2022, 7, 31), datetime.date(2023, 9, 18), datetime.date(2023, 9, 24), datetime.date(2022, 7, 30), datetime.date(2023, 10, 18), datetime.date(2023, 10, 19), datetime.date(2023, 4, 23), datetime.date(2023, 4, 28), datetime.date(2023, 5, 22), datetime.date(2023, 6, 2), datetime.date(2023, 10, 13), datetime.date(2023, 10, 24), datetime.date(2023, 10, 25), datetime.date(2023, 10, 6), datetime.date(2022, 6, 27), datetime.date(2022, 6, 27), datetime.date(2022, 6, 30), datetime.date(2022, 6, 30), datetime.date(2023, 10, 13), datetime.date(2023, 3, 15), datetime.date(2023, 9, 22), datetime.date(2022, 3, 29), datetime.date(2022, 4, 1), datetime.date(2023, 9, 22), datetime.date(2022, 1, 6), datetime.date(2023, 11, 21), datetime.date(2023, 12, 5), datetime.date(2023, 6, 5), datetime.date(2023, 12, 13), datetime.date(2023, 12, 20), datetime.date(2023, 12, 22), datetime.date(2023, 12, 23), datetime.date(2023, 12, 23), datetime.date(2023, 9, 21), datetime.date(2023, 12, 1), datetime.date(2024, 1, 28), datetime.date(2024, 1, 29), datetime.date(2024, 1, 31), datetime.date(2024, 2, 23), datetime.date(2024, 3, 7), datetime.date(2024, 3, 23), datetime.date(2023, 11, 11), datetime.date(2023, 11, 29), datetime.date(2022, 6, 28), datetime.date(2023, 12, 27), datetime.date(2023, 12, 6), datetime.date(2023, 12, 30), datetime.date(2023, 12, 28), datetime.date(2023, 12, 26), datetime.date(2023, 12, 2), datetime.date(2024, 2, 26), datetime.date(2023, 2, 26), datetime.date(2024, 2, 21), datetime.date(2024, 5, 20), datetime.date(2024, 5, 27), datetime.date(2024, 6, 12), datetime.date(2024, 6, 14), datetime.date(2024, 5, 29), datetime.date(2024, 6, 16), datetime.date(2024, 6, 19), datetime.date(2024, 6, 11), datetime.date(2024, 7, 1), datetime.date(2024, 6, 19), datetime.date(2024, 6, 24), datetime.date(2024, 6, 20), datetime.date(2024, 6, 11), datetime.date(2024, 7, 5), datetime.date(2024, 7, 21), datetime.date(2024, 7, 19), datetime.date(2024, 7, 27), datetime.date(2024, 7, 27), datetime.date(2024, 8, 1), datetime.date(2024, 8, 3), datetime.date(2024, 8, 4), datetime.date(2024, 8, 9), datetime.date(2024, 6, 14), datetime.date(2024, 8, 15), datetime.date(2024, 8, 30), datetime.date(2024, 7, 3), datetime.date(2023, 10, 4), datetime.date(2024, 10, 7), datetime.date(2024, 9, 29), datetime.date(2024, 9, 29), datetime.date(2024, 9, 30), datetime.date(2024, 9, 23), datetime.date(2024, 9, 27), datetime.date(2024, 9, 18), datetime.date(2024, 9, 19), datetime.date(2024, 9, 19), datetime.date(2024, 9, 24), datetime.date(2024, 9, 24), datetime.date(2024, 9, 24), datetime.date(2024, 9, 21), datetime.date(2024, 9, 19), datetime.date(2024, 9, 19), datetime.date(2024, 9, 24), datetime.date(2024, 9, 24), datetime.date(2024, 9, 24), datetime.date(2024, 10, 8), datetime.date(2024, 10, 6), datetime.date(2024, 11, 10), datetime.date(2024, 7, 5), datetime.date(2024, 8, 31), datetime.date(2024, 11, 26), datetime.date(2022, 6, 16), datetime.date(2024, 11, 8), datetime.date(2024, 11, 5), datetime.date(2024, 10, 2), datetime.date(2024, 11, 19), datetime.date(2024, 11, 28), datetime.date(2024, 12, 5), datetime.date(2024, 12, 16), datetime.date(2024, 12, 16), datetime.date(2024, 12, 8), datetime.date(2024, 12, 2), datetime.date(2024, 12, 4), datetime.date(2024, 12, 7), datetime.date(2024, 12, 7), datetime.date(2024, 12, 1), datetime.date(2024, 1, 9), datetime.date(2024, 6, 30), datetime.date(2024, 12, 28), datetime.date(2025, 1, 11), datetime.date(2025, 1, 9), datetime.date(2024, 8, 19), datetime.date(2025, 1, 20), datetime.date(2024, 12, 9), datetime.date(2025, 1, 15), datetime.date(2025, 1, 24), datetime.date(2025, 2, 12), datetime.date(2025, 2, 3), datetime.date(2025, 2, 10), datetime.date(2025, 1, 29), datetime.date(2025, 2, 14), datetime.date(2025, 1, 10), datetime.date(2024, 11, 22), datetime.date(2024, 11, 24), datetime.date(2025, 2, 26), datetime.date(2025, 3, 11), datetime.date(2025, 3, 1), datetime.date(2025, 2, 13), datetime.date(2025, 2, 14), datetime.date(2025, 2, 14), datetime.date(2025, 2, 10), datetime.date(2025, 2, 13), datetime.date(2025, 3, 2), datetime.date(2024, 8, 13), datetime.date(2023, 11, 28), datetime.date(2021, 12, 3), datetime.date(2021, 11, 26), datetime.date(2021, 11, 30), datetime.date(2021, 11, 25), datetime.date(2021, 11, 18), datetime.date(2021, 11, 29), datetime.date(2021, 11, 29), datetime.date(2021, 11, 25), datetime.date(2021, 11, 30), datetime.date(2021, 12, 13), datetime.date(2021, 12, 2), datetime.date(2021, 12, 7), datetime.date(2021, 12, 1), datetime.date(2021, 12, 6), datetime.date(2021, 12, 6), datetime.date(2021, 12, 14), datetime.date(2021, 12, 10), datetime.date(2021, 12, 16), datetime.date(2021, 12, 15), datetime.date(2021, 12, 18), datetime.date(2021, 12, 10), datetime.date(2021, 12, 15), datetime.date(2021, 12, 2), datetime.date(2021, 12, 21), datetime.date(2021, 12, 28), datetime.date(2021, 12, 14), datetime.date(2021, 11, 23), datetime.date(2021, 11, 29), datetime.date(2021, 11, 30), datetime.date(2021, 11, 30), datetime.date(2021, 12, 1), datetime.date(2021, 12, 13), datetime.date(2022, 1, 2), datetime.date(2021, 12, 17), datetime.date(2022, 1, 4), datetime.date(2021, 12, 15), datetime.date(2021, 11, 24), datetime.date(2021, 11, 24), datetime.date(2021, 11, 22), datetime.date(2021, 12, 14), datetime.date(2022, 1, 3), datetime.date(2022, 1, 14), datetime.date(2022, 1, 14), datetime.date(2021, 12, 30), datetime.date(2021, 12, 31), datetime.date(2022, 1, 1), datetime.date(2022, 1, 1), datetime.date(2021, 12, 30), datetime.date(2021, 12, 31), datetime.date(2021, 12, 31), datetime.date(2022, 1, 4), datetime.date(2021, 11, 30), datetime.date(2021, 12, 2), datetime.date(2021, 12, 3), datetime.date(2021, 12, 2), datetime.date(2021, 12, 3), datetime.date(2021, 12, 14), datetime.date(2021, 12, 20), datetime.date(2022, 1, 7), datetime.date(2021, 12, 2), datetime.date(2021, 12, 3), datetime.date(2021, 12, 3), datetime.date(2021, 12, 2), datetime.date(2022, 1, 3), datetime.date(2021, 12, 26), datetime.date(2022, 1, 8), datetime.date(2022, 1, 16), datetime.date(2022, 1, 20), datetime.date(2022, 1, 4), datetime.date(2022, 1, 5), datetime.date(2022, 1, 5), datetime.date(2022, 1, 4), datetime.date(2022, 1, 5), datetime.date(2022, 1, 6), datetime.date(2022, 1, 6), datetime.date(2022, 1, 4), datetime.date(2022, 1, 6), datetime.date(2022, 1, 6), datetime.date(2022, 1, 6), datetime.date(2022, 1, 3), datetime.date(2022, 1, 5), datetime.date(2022, 1, 6), datetime.date(2022, 1, 9), datetime.date(2022, 1, 11), datetime.date(2022, 1, 4), datetime.date(2022, 1, 3), datetime.date(2022, 1, 3), datetime.date(2022, 1, 3), datetime.date(2022, 1, 6), datetime.date(2022, 1, 6), datetime.date(2022, 1, 11), datetime.date(2022, 1, 13), datetime.date(2022, 1, 6), datetime.date(2022, 1, 14), datetime.date(2022, 1, 12), datetime.date(2022, 1, 9), datetime.date(2021, 12, 7), datetime.date(2021, 12, 7), datetime.date(2021, 12, 8), datetime.date(2021, 12, 11), datetime.date(2022, 1, 8), datetime.date(2022, 1, 8), datetime.date(2022, 1, 11), datetime.date(2022, 1, 15), datetime.date(2022, 1, 17), datetime.date(2022, 1, 14), datetime.date(2022, 1, 3), datetime.date(2022, 1, 3), datetime.date(2022, 1, 3), datetime.date(2022, 1, 4), datetime.date(2022, 1, 4), datetime.date(2022, 1, 4), datetime.date(2022, 1, 4), datetime.date(2022, 1, 5), datetime.date(2022, 1, 6), datetime.date(2022, 1, 7), datetime.date(2022, 1, 7), datetime.date(2022, 1, 7), datetime.date(2022, 1, 8), datetime.date(2022, 1, 7), datetime.date(2022, 1, 8), datetime.date(2022, 1, 9), datetime.date(2022, 1, 9), datetime.date(2022, 1, 9), datetime.date(2022, 1, 9), datetime.date(2022, 1, 7), datetime.date(2022, 1, 3), datetime.date(2022, 1, 7), datetime.date(2022, 1, 8), datetime.date(2022, 1, 5), datetime.date(2022, 1, 12), datetime.date(2022, 1, 15), datetime.date(2022, 1, 15), datetime.date(2022, 1, 11), datetime.date(2022, 1, 5), datetime.date(2022, 1, 7), datetime.date(2022, 1, 25), datetime.date(2022, 1, 22), datetime.date(2022, 1, 24), datetime.date(2022, 1, 17), datetime.date(2022, 1, 11), datetime.date(2022, 1, 14), datetime.date(2022, 1, 14), datetime.date(2022, 1, 14), datetime.date(2022, 1, 14), datetime.date(2022, 1, 15), datetime.date(2022, 1, 15), datetime.date(2022, 1, 14), datetime.date(2022, 1, 14), datetime.date(2022, 1, 15), datetime.date(2022, 1, 15), datetime.date(2022, 1, 15), datetime.date(2022, 1, 15), datetime.date(2022, 1, 15), datetime.date(2022, 1, 16), datetime.date(2022, 1, 14), datetime.date(2022, 1, 11), datetime.date(2022, 1, 14), datetime.date(2022, 1, 14), datetime.date(2022, 1, 10), datetime.date(2022, 1, 10), datetime.date(2022, 1, 10), datetime.date(2022, 1, 10), datetime.date(2022, 1, 10), datetime.date(2022, 1, 10), datetime.date(2022, 1, 11), datetime.date(2022, 1, 11), datetime.date(2022, 1, 11), datetime.date(2022, 1, 10), datetime.date(2022, 1, 11), datetime.date(2022, 1, 12), datetime.date(2022, 1, 12), datetime.date(2022, 1, 13), datetime.date(2022, 1, 14), datetime.date(2022, 1, 13), datetime.date(2022, 1, 14), datetime.date(2022, 1, 14), datetime.date(2022, 1, 14), datetime.date(2022, 1, 27), datetime.date(2022, 1, 20), datetime.date(2022, 1, 23), datetime.date(2022, 1, 20), datetime.date(2022, 1, 13), datetime.date(2022, 1, 15), datetime.date(2022, 1, 16), datetime.date(2022, 1, 10), datetime.date(2022, 1, 16), datetime.date(2022, 1, 8), datetime.date(2022, 1, 8), datetime.date(2021, 12, 17), datetime.date(2022, 1, 13), datetime.date(2022, 1, 25), datetime.date(2022, 1, 24), datetime.date(2022, 2, 1), datetime.date(2021, 12, 8), datetime.date(2021, 12, 8), datetime.date(2021, 12, 16), datetime.date(2022, 1, 11), datetime.date(2022, 1, 18), datetime.date(2022, 1, 11), datetime.date(2022, 1, 11), datetime.date(2022, 1, 13), datetime.date(2021, 12, 4), datetime.date(2021, 12, 15), datetime.date(2022, 1, 3), datetime.date(2022, 1, 3), datetime.date(2022, 1, 24), datetime.date(2022, 1, 19), datetime.date(2022, 1, 19), datetime.date(2022, 1, 20), datetime.date(2022, 1, 21), datetime.date(2022, 1, 17), datetime.date(2022, 1, 18), datetime.date(2022, 1, 17), datetime.date(2022, 1, 18), datetime.date(2022, 1, 18), datetime.date(2022, 1, 17), datetime.date(2022, 1, 17), datetime.date(2022, 1, 19), datetime.date(2022, 1, 19), datetime.date(2022, 1, 19), datetime.date(2022, 1, 19), datetime.date(2022, 1, 19), datetime.date(2022, 1, 19), datetime.date(2022, 1, 20), datetime.date(2022, 1, 20), datetime.date(2022, 1, 20), datetime.date(2022, 1, 20), datetime.date(2022, 1, 19), datetime.date(2022, 1, 21), datetime.date(2022, 1, 21), datetime.date(2022, 1, 21), datetime.date(2022, 1, 21), datetime.date(2022, 1, 22), datetime.date(2022, 1, 22), datetime.date(2022, 1, 23), datetime.date(2022, 1, 18), datetime.date(2022, 1, 22), datetime.date(2022, 1, 21), datetime.date(2022, 1, 22), datetime.date(2022, 1, 19), datetime.date(2022, 1, 19), datetime.date(2022, 1, 21), datetime.date(2022, 1, 19), datetime.date(2022, 1, 18), datetime.date(2022, 1, 19), datetime.date(2022, 1, 20), datetime.date(2022, 1, 21), datetime.date(2022, 1, 21), datetime.date(2022, 1, 18), datetime.date(2022, 1, 20), datetime.date(2022, 1, 21), datetime.date(2022, 1, 18), datetime.date(2022, 1, 19), datetime.date(2022, 1, 20), datetime.date(2022, 2, 3), datetime.date(2022, 1, 20), datetime.date(2022, 1, 31), datetime.date(2022, 1, 25), datetime.date(2022, 1, 26), datetime.date(2022, 1, 12), datetime.date(2022, 1, 17), datetime.date(2022, 1, 31), datetime.date(2022, 1, 24), datetime.date(2022, 1, 27), datetime.date(2021, 12, 20), datetime.date(2021, 12, 29), datetime.date(2022, 1, 18), datetime.date(2022, 1, 28), datetime.date(2022, 1, 23), datetime.date(2022, 1, 25), datetime.date(2022, 1, 24), datetime.date(2022, 1, 25), datetime.date(2022, 1, 23), datetime.date(2022, 1, 25), datetime.date(2021, 12, 4), datetime.date(2022, 1, 19), datetime.date(2022, 1, 20), datetime.date(2022, 1, 18), datetime.date(2022, 1, 17), datetime.date(2022, 1, 17), datetime.date(2022, 1, 17), datetime.date(2022, 1, 17), datetime.date(2022, 1, 17), datetime.date(2022, 1, 17), datetime.date(2022, 1, 17), datetime.date(2022, 1, 17), datetime.date(2022, 1, 15), datetime.date(2022, 1, 17), datetime.date(2022, 1, 17), datetime.date(2022, 1, 17), datetime.date(2022, 1, 25), datetime.date(2022, 1, 25), datetime.date(2022, 1, 24), datetime.date(2022, 1, 25), datetime.date(2022, 1, 26), datetime.date(2022, 1, 26), datetime.date(2022, 1, 26), datetime.date(2022, 1, 27), datetime.date(2022, 1, 17), datetime.date(2022, 1, 22), datetime.date(2022, 1, 22), datetime.date(2022, 1, 22), datetime.date(2022, 1, 24), datetime.date(2022, 1, 24), datetime.date(2022, 1, 23), datetime.date(2022, 1, 26), datetime.date(2022, 1, 20), datetime.date(2022, 1, 22), datetime.date(2022, 1, 26), datetime.date(2022, 1, 28), datetime.date(2022, 1, 30), datetime.date(2022, 1, 20), datetime.date(2022, 1, 24), datetime.date(2022, 1, 24), datetime.date(2022, 1, 25), datetime.date(2022, 1, 25), datetime.date(2022, 2, 2), datetime.date(2022, 1, 12), datetime.date(2022, 1, 27), datetime.date(2022, 1, 31), datetime.date(2022, 1, 27), datetime.date(2022, 1, 27), datetime.date(2022, 1, 27), datetime.date(2022, 1, 28), datetime.date(2022, 1, 28), datetime.date(2022, 1, 28), datetime.date(2022, 1, 27), datetime.date(2022, 1, 29), datetime.date(2022, 1, 28), datetime.date(2022, 1, 30), datetime.date(2022, 1, 29), datetime.date(2022, 1, 30), datetime.date(2022, 1, 29), datetime.date(2022, 1, 30), datetime.date(2022, 1, 27), datetime.date(2022, 1, 27), datetime.date(2022, 1, 30), datetime.date(2022, 1, 26), datetime.date(2022, 1, 26), datetime.date(2022, 1, 24), datetime.date(2022, 1, 27), datetime.date(2022, 1, 27), datetime.date(2021, 12, 15), datetime.date(2021, 12, 15), datetime.date(2021, 12, 29), datetime.date(2021, 12, 26), datetime.date(2022, 1, 15), datetime.date(2021, 12, 3), datetime.date(2021, 12, 25), datetime.date(2021, 12, 19), datetime.date(2022, 1, 4), datetime.date(2021, 12, 23), datetime.date(2022, 1, 4), datetime.date(2022, 2, 1), datetime.date(2022, 2, 1), datetime.date(2022, 2, 3), datetime.date(2022, 1, 26), datetime.date(2022, 2, 11), datetime.date(2022, 2, 10)]
```

In [10]:

```
# grouping dates into months, year

from collections import defaultdict

date_groups = defaultdict(list)

for date in collection_dates:
    # date = datetime.strptime(date_str, '%Y-%-%d')
    month_year_key = date.strftime('%Y-%m')    # Key Format: '2002-01'
    date_groups[month_year_key].append(date)

# Print the grouped dates
for month_year, dates_in_group in date_groups.items():
    print(f'Dates in {month_year}:{dates_in_group}')
```

```
Dates in 2022-02:[datetime.date(2022, 2, 3), datetime.date(2022, 2, 8), datetime.date(2022, 2, 3), datetime.date(2022, 2, 7), datetime.date(2022, 2, 6), datetime.date(2022, 2, 13), datetime.date(2022, 2, 12), datetime.date(2022, 2, 2), datetime.date(2022, 2, 4), datetime.date(2022, 2, 6), datetime.date(2022, 2, 2), datetime.date(2022, 2, 15), datetime.date(2022, 2, 1), datetime.date(2022, 2, 2), datetime.date(2022, 2, 4), datetime.date(2022, 2, 4), datetime.date(2022, 2, 3), datetime.date(2022, 2, 3), datetime.date(2022, 2, 4), datetime.date(2022, 2, 3), datetime.date(2022, 2, 3), datetime.date(2022, 2, 1), datetime.date(2022, 2, 1), datetime.date(2022, 2, 7), datetime.date(2022, 2, 8), datetime.date(2022, 2, 10), datetime.date(2022, 2, 12), datetime.date(2022, 2, 14), datetime.date(2022, 2, 21), datetime.date(2022, 2, 9), datetime.date(2022, 2, 9), datetime.date(2022, 2, 13), datetime.date(2022, 2, 18), datetime.date(2022, 2, 7), datetime.date(2022, 2, 6), datetime.date(2022, 2, 14), datetime.date(2022, 2, 14), datetime.date(2022, 2, 11), datetime.date(2022, 2, 1), datetime.date(2022, 2, 5), datetime.date(2022, 2, 28), datetime.date(2022, 2, 15), datetime.date(2022, 2, 1), datetime.date(2022, 2, 9), datetime.date(2022, 2, 3), datetime.date(2022, 2, 14), datetime.date(2022, 2, 16), datetime.date(2022, 2, 28), datetime.date(2022, 2, 8), datetime.date(2022, 2, 15), datetime.date(2022, 2, 17), datetime.date(2022, 2, 18), datetime.date(2022, 2, 8), datetime.date(2022, 2, 28), datetime.date(2022, 2, 15), datetime.date(2022, 2, 16), datetime.date(2022, 2, 16), datetime.date(2022, 2, 13), datetime.date(2022, 2, 22), datetime.date(2022, 2, 17), datetime.date(2022, 2, 20), datetime.date(2022, 2, 17), datetime.date(2022, 2, 17), datetime.date(2022, 2, 4), datetime.date(2022, 2, 9), datetime.date(2022, 2, 21), datetime.date(2022, 2, 22), datetime.date(2022, 2, 22), datetime.date(2022, 2, 22), datetime.date(2022, 2, 21), datetime.date(2022, 2, 24), datetime.date(2022, 2, 21), datetime.date(2022, 2, 20), datetime.date(2022, 2, 18), datetime.date(2022, 2, 20), datetime.date(2022, 2, 4), datetime.date(2022, 2, 21), datetime.date(2022, 2, 14), datetime.date(2022, 2, 1), datetime.date(2022, 2, 15), datetime.date(2022, 2, 7), datetime.date(2022, 2, 24), datetime.date(2022, 2, 14), datetime.date(2022, 2, 4), datetime.date(2022, 2, 1), datetime.date(2022, 2, 3), datetime.date(2022, 2, 2), datetime.date(2022, 2, 1), datetime.date(2022, 2, 1), datetime.date(2022, 2, 3), datetime.date(2022, 2, 11), datetime.date(2022, 2, 10)]
Dates in 2022-01:[datetime.date(2022, 1, 29), datetime.date(2022, 1, 27), datetime.date(2022, 1, 28), datetime.date(2022, 1, 29), datetime.date(2022, 1, 28), datetime.date(2022, 1, 31), datetime.date(2022, 1, 31), datetime.date(2022, 1, 31), datetime.date(2022, 1, 2), datetime.date(2022, 1, 7), datetime.date(2022, 1, 11), datetime.date(2022, 1, 8), datetime.date(2022, 1, 31), datetime.date(2022, 1, 31), datetime.date(2022, 1, 27), datetime.date(2022, 1, 22), datetime.date(2022, 1, 21), datetime.date(2022, 1, 22), datetime.date(2022, 1, 5), datetime.date(2022, 1, 18), datetime.date(2022, 1, 28), datetime.date(2022, 1, 28), datetime.date(2022, 1, 28), datetime.date(2022, 1, 20), datetime.date(2022, 1, 5), datetime.date(2022, 1, 27), datetime.date(2022, 1, 28), datetime.date(2022, 1, 30), datetime.date(2022, 1, 30), datetime.date(2022, 1, 31), datetime.date(2022, 1, 23), datetime.date(2022, 1, 30), datetime.date(2022, 1, 19), datetime.date(2022, 1, 25), datetime.date(2022, 1, 27), datetime.date(2022, 1, 30), datetime.date(2022, 1, 29), datetime.date(2022, 1, 30), datetime.date(2022, 1, 31), datetime.date(2022, 1, 19), datetime.date(2022, 1, 5), datetime.date(2022, 1, 6), datetime.date(2022, 1, 2), datetime.date(2022, 1, 4), datetime.date(2022, 1, 3), datetime.date(2022, 1, 14), datetime.date(2022, 1, 14), datetime.date(2022, 1, 1), datetime.date(2022, 1, 1), datetime.date(2022, 1, 4), datetime.date(2022, 1, 7), datetime.date(2022, 1, 3), datetime.date(2022, 1, 8), datetime.date(2022, 1, 16), datetime.date(2022, 1, 20), datetime.date(2022, 1, 4), datetime.date(2022, 1, 5), datetime.date(2022, 1, 5), datetime.date(2022, 1, 4), datetime.date(2022, 1, 5), datetime.date(2022, 1, 6), datetime.date(2022, 1, 6), datetime.date(2022, 1, 4), datetime.date(2022, 1, 6), datetime.date(2022, 1, 6), datetime.date(2022, 1, 6), datetime.date(2022, 1, 3), datetime.date(2022, 1, 5), datetime.date(2022, 1, 6), datetime.date(2022, 1, 9), datetime.date(2022, 1, 11), datetime.date(2022, 1, 4), datetime.date(2022, 1, 3), datetime.date(2022, 1, 3), datetime.date(2022, 1, 3), datetime.date(2022, 1, 6), datetime.date(2022, 1, 6), datetime.date(2022, 1, 11), datetime.date(2022, 1, 13), datetime.date(2022, 1, 6), datetime.date(2022, 1, 14), datetime.date(2022, 1, 12), datetime.date(2022, 1, 9), datetime.date(2022, 1, 8), datetime.date(2022, 1, 8), datetime.date(2022, 1, 11), datetime.date(2022, 1, 15), datetime.date(2022, 1, 17), datetime.date(2022, 1, 14), datetime.date(2022, 1, 3), datetime.date(2022, 1, 3), datetime.date(2022, 1, 3), datetime.date(2022, 1, 4), datetime.date(2022, 1, 4), datetime.date(2022, 1, 4), datetime.date(2022, 1, 4), datetime.date(2022, 1, 5), datetime.date(2022, 1, 6), datetime.date(2022, 1, 7), datetime.date(2022, 1, 7), datetime.date(2022, 1, 7), datetime.date(2022, 1, 8), datetime.date(2022, 1, 7), datetime.date(2022, 1, 8), datetime.date(2022, 1, 9), datetime.date(2022, 1, 9), datetime.date(2022, 1, 9), datetime.date(2022, 1, 9), datetime.date(2022, 1, 7), datetime.date(2022, 1, 3), datetime.date(2022, 1, 7), datetime.date(2022, 1, 8), datetime.date(2022, 1, 5), datetime.date(2022, 1, 12), datetime.date(2022, 1, 15), datetime.date(2022, 1, 15), datetime.date(2022, 1, 11), datetime.date(2022, 1, 5), datetime.date(2022, 1, 7), datetime.date(2022, 1, 25), datetime.date(2022, 1, 22), datetime.date(2022, 1, 24), datetime.date(2022, 1, 17), datetime.date(2022, 1, 11), datetime.date(2022, 1, 14), datetime.date(2022, 1, 14), datetime.date(2022, 1, 14), datetime.date(2022, 1, 14), datetime.date(2022, 1, 15), datetime.date(2022, 1, 15), datetime.date(2022, 1, 14), datetime.date(2022, 1, 14), datetime.date(2022, 1, 15), datetime.date(2022, 1, 15), datetime.date(2022, 1, 15), datetime.date(2022, 1, 15), datetime.date(2022, 1, 15), datetime.date(2022, 1, 16), datetime.date(2022, 1, 14), datetime.date(2022, 1, 11), datetime.date(2022, 1, 14), datetime.date(2022, 1, 14), datetime.date(2022, 1, 10), datetime.date(2022, 1, 10), datetime.date(2022, 1, 10), datetime.date(2022, 1, 10), datetime.date(2022, 1, 10), datetime.date(2022, 1, 10), datetime.date(2022, 1, 11), datetime.date(2022, 1, 11), datetime.date(2022, 1, 11), datetime.date(2022, 1, 10), datetime.date(2022, 1, 11), datetime.date(2022, 1, 12), datetime.date(2022, 1, 12), datetime.date(2022, 1, 13), datetime.date(2022, 1, 14), datetime.date(2022, 1, 13), datetime.date(2022, 1, 14), datetime.date(2022, 1, 14), datetime.date(2022, 1, 14), datetime.date(2022, 1, 27), datetime.date(2022, 1, 20), datetime.date(2022, 1, 23), datetime.date(2022, 1, 20), datetime.date(2022, 1, 13), datetime.date(2022, 1, 15), datetime.date(2022, 1, 16), datetime.date(2022, 1, 10), datetime.date(2022, 1, 16), datetime.date(2022, 1, 8), datetime.date(2022, 1, 8), datetime.date(2022, 1, 13), datetime.date(2022, 1, 25), datetime.date(2022, 1, 24), datetime.date(2022, 1, 11), datetime.date(2022, 1, 18), datetime.date(2022, 1, 11), datetime.date(2022, 1, 11), datetime.date(2022, 1, 13), datetime.date(2022, 1, 3), datetime.date(2022, 1, 3), datetime.date(2022, 1, 24), datetime.date(2022, 1, 19), datetime.date(2022, 1, 19), datetime.date(2022, 1, 20), datetime.date(2022, 1, 21), datetime.date(2022, 1, 17), datetime.date(2022, 1, 18), datetime.date(2022, 1, 17), datetime.date(2022, 1, 18), datetime.date(2022, 1, 18), datetime.date(2022, 1, 17), datetime.date(2022, 1, 17), datetime.date(2022, 1, 19), datetime.date(2022, 1, 19), datetime.date(2022, 1, 19), datetime.date(2022, 1, 19), datetime.date(2022, 1, 19), datetime.date(2022, 1, 19), datetime.date(2022, 1, 20), datetime.date(2022, 1, 20), datetime.date(2022, 1, 20), datetime.date(2022, 1, 20), datetime.date(2022, 1, 19), datetime.date(2022, 1, 21), datetime.date(2022, 1, 21), datetime.date(2022, 1, 21), datetime.date(2022, 1, 21), datetime.date(2022, 1, 22), datetime.date(2022, 1, 22), datetime.date(2022, 1, 23), datetime.date(2022, 1, 18), datetime.date(2022, 1, 22), datetime.date(2022, 1, 21), datetime.date(2022, 1, 22), datetime.date(2022, 1, 19), datetime.date(2022, 1, 19), datetime.date(2022, 1, 21), datetime.date(2022, 1, 19), datetime.date(2022, 1, 18), datetime.date(2022, 1, 19), datetime.date(2022, 1, 20), datetime.date(2022, 1, 21), datetime.date(2022, 1, 21), datetime.date(2022, 1, 18), datetime.date(2022, 1, 20), datetime.date(2022, 1, 21), datetime.date(2022, 1, 18), datetime.date(2022, 1, 19), datetime.date(2022, 1, 20), datetime.date(2022, 1, 20), datetime.date(2022, 1, 31), datetime.date(2022, 1, 25), datetime.date(2022, 1, 26), datetime.date(2022, 1, 12), datetime.date(2022, 1, 17), datetime.date(2022, 1, 31), datetime.date(2022, 1, 24), datetime.date(2022, 1, 27), datetime.date(2022, 1, 18), datetime.date(2022, 1, 28), datetime.date(2022, 1, 23), datetime.date(2022, 1, 25), datetime.date(2022, 1, 24), datetime.date(2022, 1, 25), datetime.date(2022, 1, 23), datetime.date(2022, 1, 25), datetime.date(2022, 1, 19), datetime.date(2022, 1, 20), datetime.date(2022, 1, 18), datetime.date(2022, 1, 17), datetime.date(2022, 1, 17), datetime.date(2022, 1, 17), datetime.date(2022, 1, 17), datetime.date(2022, 1, 17), datetime.date(2022, 1, 17), datetime.date(2022, 1, 17), datetime.date(2022, 1, 17), datetime.date(2022, 1, 15), datetime.date(2022, 1, 17), datetime.date(2022, 1, 17), datetime.date(2022, 1, 17), datetime.date(2022, 1, 25), datetime.date(2022, 1, 25), datetime.date(2022, 1, 24), datetime.date(2022, 1, 25), datetime.date(2022, 1, 26), datetime.date(2022, 1, 26), datetime.date(2022, 1, 26), datetime.date(2022, 1, 27), datetime.date(2022, 1, 17), datetime.date(2022, 1, 22), datetime.date(2022, 1, 22), datetime.date(2022, 1, 22), datetime.date(2022, 1, 24), datetime.date(2022, 1, 24), datetime.date(2022, 1, 23), datetime.date(2022, 1, 26), datetime.date(2022, 1, 20), datetime.date(2022, 1, 22), datetime.date(2022, 1, 26), datetime.date(2022, 1, 28), datetime.date(2022, 1, 30), datetime.date(2022, 1, 20), datetime.date(2022, 1, 24), datetime.date(2022, 1, 24), datetime.date(2022, 1, 25), datetime.date(2022, 1, 25), datetime.date(2022, 1, 12), datetime.date(2022, 1, 27), datetime.date(2022, 1, 31), datetime.date(2022, 1, 27), datetime.date(2022, 1, 27), datetime.date(2022, 1, 27), datetime.date(2022, 1, 28), datetime.date(2022, 1, 28), datetime.date(2022, 1, 28), datetime.date(2022, 1, 27), datetime.date(2022, 1, 29), datetime.date(2022, 1, 28), datetime.date(2022, 1, 30), datetime.date(2022, 1, 29), datetime.date(2022, 1, 30), datetime.date(2022, 1, 29), datetime.date(2022, 1, 30), datetime.date(2022, 1, 27), datetime.date(2022, 1, 27), datetime.date(2022, 1, 30), datetime.date(2022, 1, 26), datetime.date(2022, 1, 26), datetime.date(2022, 1, 24), datetime.date(2022, 1, 27), datetime.date(2022, 1, 27), datetime.date(2022, 1, 15), datetime.date(2022, 1, 4), datetime.date(2022, 1, 4), datetime.date(2022, 1, 26)]
Dates in 2021-12:[datetime.date(2021, 12, 4), datetime.date(2021, 12, 8), datetime.date(2021, 12, 8), datetime.date(2021, 12, 6), datetime.date(2021, 12, 24), datetime.date(2021, 12, 1), datetime.date(2021, 12, 7), datetime.date(2021, 12, 9), datetime.date(2021, 12, 9), datetime.date(2021, 12, 6), datetime.date(2021, 12, 28), datetime.date(2021, 12, 31), datetime.date(2021, 12, 31), datetime.date(2021, 12, 31), datetime.date(2021, 12, 31), datetime.date(2021, 12, 20), datetime.date(2021, 12, 20), datetime.date(2021, 12, 20), datetime.date(2021, 12, 1), datetime.date(2021, 12, 25), datetime.date(2021, 12, 6), datetime.date(2021, 12, 24), datetime.date(2021, 12, 2), datetime.date(2021, 12, 6), datetime.date(2021, 12, 9), datetime.date(2021, 12, 9), datetime.date(2021, 12, 9), datetime.date(2021, 12, 10), datetime.date(2021, 12, 10), datetime.date(2021, 12, 10), datetime.date(2021, 12, 6), datetime.date(2021, 12, 6), datetime.date(2021, 12, 10), datetime.date(2021, 12, 10), datetime.date(2021, 12, 6), datetime.date(2021, 12, 6), datetime.date(2021, 12, 6), datetime.date(2021, 12, 7), datetime.date(2021, 12, 13), datetime.date(2021, 12, 21), datetime.date(2021, 12, 8), datetime.date(2021, 12, 8), datetime.date(2021, 12, 6), datetime.date(2021, 12, 10), datetime.date(2021, 12, 10), datetime.date(2021, 12, 13), datetime.date(2021, 12, 7), datetime.date(2021, 12, 15), datetime.date(2021, 12, 3), datetime.date(2021, 12, 4), datetime.date(2021, 12, 3), datetime.date(2021, 12, 13), datetime.date(2021, 12, 2), datetime.date(2021, 12, 7), datetime.date(2021, 12, 1), datetime.date(2021, 12, 6), datetime.date(2021, 12, 6), datetime.date(2021, 12, 14), datetime.date(2021, 12, 10), datetime.date(2021, 12, 16), datetime.date(2021, 12, 15), datetime.date(2021, 12, 18), datetime.date(2021, 12, 10), datetime.date(2021, 12, 15), datetime.date(2021, 12, 2), datetime.date(2021, 12, 21), datetime.date(2021, 12, 28), datetime.date(2021, 12, 14), datetime.date(2021, 12, 1), datetime.date(2021, 12, 13), datetime.date(2021, 12, 17), datetime.date(2021, 12, 15), datetime.date(2021, 12, 14), datetime.date(2021, 12, 30), datetime.date(2021, 12, 31), datetime.date(2021, 12, 30), datetime.date(2021, 12, 31), datetime.date(2021, 12, 31), datetime.date(2021, 12, 2), datetime.date(2021, 12, 3), datetime.date(2021, 12, 2), datetime.date(2021, 12, 3), datetime.date(2021, 12, 14), datetime.date(2021, 12, 20), datetime.date(2021, 12, 2), datetime.date(2021, 12, 3), datetime.date(2021, 12, 3), datetime.date(2021, 12, 2), datetime.date(2021, 12, 26), datetime.date(2021, 12, 7), datetime.date(2021, 12, 7), datetime.date(2021, 12, 8), datetime.date(2021, 12, 11), datetime.date(2021, 12, 17), datetime.date(2021, 12, 8), datetime.date(2021, 12, 8), datetime.date(2021, 12, 16), datetime.date(2021, 12, 4), datetime.date(2021, 12, 15), datetime.date(2021, 12, 20), datetime.date(2021, 12, 29), datetime.date(2021, 12, 4), datetime.date(2021, 12, 15), datetime.date(2021, 12, 15), datetime.date(2021, 12, 29), datetime.date(2021, 12, 26), datetime.date(2021, 12, 3), datetime.date(2021, 12, 25), datetime.date(2021, 12, 19), datetime.date(2021, 12, 23)]
Dates in 2022-03:[datetime.date(2022, 3, 7), datetime.date(2022, 3, 7), datetime.date(2022, 3, 7), datetime.date(2022, 3, 15), datetime.date(2022, 3, 4), datetime.date(2022, 3, 16), datetime.date(2022, 3, 15), datetime.date(2022, 3, 22), datetime.date(2022, 3, 23), datetime.date(2022, 3, 23), datetime.date(2022, 3, 23), datetime.date(2022, 3, 25), datetime.date(2022, 3, 26), datetime.date(2022, 3, 10), datetime.date(2022, 3, 16), datetime.date(2022, 3, 23), datetime.date(2022, 3, 14), datetime.date(2022, 3, 11), datetime.date(2022, 3, 16), datetime.date(2022, 3, 29)]
Dates in 2022-04:[datetime.date(2022, 4, 1), datetime.date(2022, 4, 17), datetime.date(2022, 4, 16), datetime.date(2022, 4, 21), datetime.date(2022, 4, 20), datetime.date(2022, 4, 15), datetime.date(2022, 4, 16), datetime.date(2022, 4, 7), datetime.date(2022, 4, 20), datetime.date(2022, 4, 23), datetime.date(2022, 4, 12), datetime.date(2022, 4, 28), datetime.date(2022, 4, 25), datetime.date(2022, 4, 25), datetime.date(2022, 4, 30), datetime.date(2022, 4, 19), datetime.date(2022, 4, 11), datetime.date(2022, 4, 7), datetime.date(2022, 4, 19), datetime.date(2022, 4, 11), datetime.date(2022, 4, 6), datetime.date(2022, 4, 19), datetime.date(2022, 4, 23), datetime.date(2022, 4, 28), datetime.date(2022, 4, 29), datetime.date(2022, 4, 27), datetime.date(2022, 4, 29), datetime.date(2022, 4, 23), datetime.date(2022, 4, 13), datetime.date(2022, 4, 1)]
Dates in 2022-05:[datetime.date(2022, 5, 1), datetime.date(2022, 5, 6), datetime.date(2022, 5, 17), datetime.date(2022, 5, 24), datetime.date(2022, 5, 27), datetime.date(2022, 5, 28), datetime.date(2022, 5, 11), datetime.date(2022, 5, 11), datetime.date(2022, 5, 31), datetime.date(2022, 5, 19), datetime.date(2022, 5, 2), datetime.date(2022, 5, 24), datetime.date(2022, 5, 5), datetime.date(2022, 5, 7), datetime.date(2022, 5, 7), datetime.date(2022, 5, 10), datetime.date(2022, 5, 27), datetime.date(2022, 5, 30), datetime.date(2022, 5, 25), datetime.date(2022, 5, 24), datetime.date(2022, 5, 8), datetime.date(2022, 5, 23), datetime.date(2022, 5, 20), datetime.date(2022, 5, 27), datetime.date(2022, 5, 12), datetime.date(2022, 5, 24), datetime.date(2022, 5, 9), datetime.date(2022, 5, 26)]
Dates in 2022-06:[datetime.date(2022, 6, 7), datetime.date(2022, 6, 6), datetime.date(2022, 6, 6), datetime.date(2022, 6, 8), datetime.date(2022, 6, 16), datetime.date(2022, 6, 16), datetime.date(2022, 6, 17), datetime.date(2022, 6, 19), datetime.date(2022, 6, 14), datetime.date(2022, 6, 16), datetime.date(2022, 6, 16), datetime.date(2022, 6, 16), datetime.date(2022, 6, 16), datetime.date(2022, 6, 16), datetime.date(2022, 6, 16), datetime.date(2022, 6, 13), datetime.date(2022, 6, 20), datetime.date(2022, 6, 29), datetime.date(2022, 6, 12), datetime.date(2022, 6, 16), datetime.date(2022, 6, 25), datetime.date(2022, 6, 14), datetime.date(2022, 6, 26), datetime.date(2022, 6, 27), datetime.date(2022, 6, 17), datetime.date(2022, 6, 27), datetime.date(2022, 6, 26), datetime.date(2022, 6, 29), datetime.date(2022, 6, 7), datetime.date(2022, 6, 14), datetime.date(2022, 6, 23), datetime.date(2022, 6, 23), datetime.date(2022, 6, 16), datetime.date(2022, 6, 30), datetime.date(2022, 6, 22), datetime.date(2022, 6, 27), datetime.date(2022, 6, 27), datetime.date(2022, 6, 30), datetime.date(2022, 6, 30), datetime.date(2022, 6, 28), datetime.date(2022, 6, 16)]
Dates in 2022-07:[datetime.date(2022, 7, 1), datetime.date(2022, 7, 4), datetime.date(2022, 7, 8), datetime.date(2022, 7, 2), datetime.date(2022, 7, 4), datetime.date(2022, 7, 8), datetime.date(2022, 7, 4), datetime.date(2022, 7, 4), datetime.date(2022, 7, 5), datetime.date(2022, 7, 5), datetime.date(2022, 7, 12), datetime.date(2022, 7, 13), datetime.date(2022, 7, 13), datetime.date(2022, 7, 2), datetime.date(2022, 7, 6), datetime.date(2022, 7, 7), datetime.date(2022, 7, 8), datetime.date(2022, 7, 10), datetime.date(2022, 7, 10), datetime.date(2022, 7, 6), datetime.date(2022, 7, 18), datetime.date(2022, 7, 24), datetime.date(2022, 7, 24), datetime.date(2022, 7, 25), datetime.date(2022, 7, 21), datetime.date(2022, 7, 25), datetime.date(2022, 7, 18), datetime.date(2022, 7, 22), datetime.date(2022, 7, 25), datetime.date(2022, 7, 15), datetime.date(2022, 7, 21), datetime.date(2022, 7, 25), datetime.date(2022, 7, 30), datetime.date(2022, 7, 11), datetime.date(2022, 7, 20), datetime.date(2022, 7, 21), datetime.date(2022, 7, 20), datetime.date(2022, 7, 8), datetime.date(2022, 7, 9), datetime.date(2022, 7, 10), datetime.date(2022, 7, 29), datetime.date(2022, 7, 30), datetime.date(2022, 7, 31), datetime.date(2022, 7, 30)]
Dates in 2022-08:[datetime.date(2022, 8, 2), datetime.date(2022, 8, 13), datetime.date(2022, 8, 11), datetime.date(2022, 8, 21), datetime.date(2022, 8, 23), datetime.date(2022, 8, 24), datetime.date(2022, 8, 23), datetime.date(2022, 8, 28), datetime.date(2022, 8, 26), datetime.date(2022, 8, 1), datetime.date(2022, 8, 2), datetime.date(2022, 8, 16), datetime.date(2022, 8, 16), datetime.date(2022, 8, 7), datetime.date(2022, 8, 31), datetime.date(2022, 8, 18), datetime.date(2022, 8, 15)]
Dates in 2022-09:[datetime.date(2022, 9, 2), datetime.date(2022, 9, 1), datetime.date(2022, 9, 9), datetime.date(2022, 9, 10), datetime.date(2022, 9, 11), datetime.date(2022, 9, 4), datetime.date(2022, 9, 21), datetime.date(2022, 9, 12), datetime.date(2022, 9, 5), datetime.date(2022, 9, 8), datetime.date(2022, 9, 10), datetime.date(2022, 9, 12), datetime.date(2022, 9, 13), datetime.date(2022, 9, 13), datetime.date(2022, 9, 10), datetime.date(2022, 9, 26), datetime.date(2022, 9, 17), datetime.date(2022, 9, 7), datetime.date(2022, 9, 3)]
Dates in 2022-10:[datetime.date(2022, 10, 9), datetime.date(2022, 10, 11), datetime.date(2022, 10, 10), datetime.date(2022, 10, 18), datetime.date(2022, 10, 10), datetime.date(2022, 10, 12), datetime.date(2022, 10, 18), datetime.date(2022, 10, 19), datetime.date(2022, 10, 18), datetime.date(2022, 10, 18), datetime.date(2022, 10, 31)]
Dates in 2022-11:[datetime.date(2022, 11, 9), datetime.date(2022, 11, 9), datetime.date(2022, 11, 5), datetime.date(2022, 11, 10), datetime.date(2022, 11, 12), datetime.date(2022, 11, 11), datetime.date(2022, 11, 1), datetime.date(2022, 11, 10), datetime.date(2022, 11, 22), datetime.date(2022, 11, 13), datetime.date(2022, 11, 10), datetime.date(2022, 11, 15), datetime.date(2022, 11, 1), datetime.date(2022, 11, 2), datetime.date(2022, 11, 1), datetime.date(2022, 11, 2), datetime.date(2022, 11, 4), datetime.date(2022, 11, 5), datetime.date(2022, 11, 26), datetime.date(2022, 11, 25), datetime.date(2022, 11, 29), datetime.date(2022, 11, 24), datetime.date(2022, 11, 27), datetime.date(2022, 11, 28), datetime.date(2022, 11, 21), datetime.date(2022, 11, 4)]
Dates in 2022-12:[datetime.date(2022, 12, 7), datetime.date(2022, 12, 6), datetime.date(2022, 12, 26), datetime.date(2022, 12, 28), datetime.date(2022, 12, 4), datetime.date(2022, 12, 3)]
Dates in 2023-01:[datetime.date(2023, 1, 2), datetime.date(2023, 1, 1), datetime.date(2023, 1, 27), datetime.date(2023, 1, 23)]
Dates in 2023-02:[datetime.date(2023, 2, 6), datetime.date(2023, 2, 13), datetime.date(2023, 2, 12), datetime.date(2023, 2, 2), datetime.date(2023, 2, 28), datetime.date(2023, 2, 22), datetime.date(2023, 2, 27), datetime.date(2023, 2, 3), datetime.date(2023, 2, 26)]
Dates in 2023-03:[datetime.date(2023, 3, 4), datetime.date(2023, 3, 3), datetime.date(2023, 3, 31), datetime.date(2023, 3, 16), datetime.date(2023, 3, 14), datetime.date(2023, 3, 13), datetime.date(2023, 3, 12), datetime.date(2023, 3, 12), datetime.date(2023, 3, 20), datetime.date(2023, 3, 26), datetime.date(2023, 3, 11), datetime.date(2023, 3, 14), datetime.date(2023, 3, 29), datetime.date(2023, 3, 24), datetime.date(2023, 3, 20), datetime.date(2023, 3, 29), datetime.date(2023, 3, 15)]
Dates in 2023-04:[datetime.date(2023, 4, 2), datetime.date(2023, 4, 1), datetime.date(2023, 4, 8), datetime.date(2023, 4, 19), datetime.date(2023, 4, 4), datetime.date(2023, 4, 12), datetime.date(2023, 4, 13), datetime.date(2023, 4, 13), datetime.date(2023, 4, 17), datetime.date(2023, 4, 14), datetime.date(2023, 4, 8), datetime.date(2023, 4, 13), datetime.date(2023, 4, 15), datetime.date(2023, 4, 19), datetime.date(2023, 4, 20), datetime.date(2023, 4, 21), datetime.date(2023, 4, 21), datetime.date(2023, 4, 24), datetime.date(2023, 4, 5), datetime.date(2023, 4, 15), datetime.date(2023, 4, 11), datetime.date(2023, 4, 11), datetime.date(2023, 4, 8), datetime.date(2023, 4, 16), datetime.date(2023, 4, 24), datetime.date(2023, 4, 21), datetime.date(2023, 4, 13), datetime.date(2023, 4, 21), datetime.date(2023, 4, 25), datetime.date(2023, 4, 15), datetime.date(2023, 4, 15), datetime.date(2023, 4, 20), datetime.date(2023, 4, 26), datetime.date(2023, 4, 23), datetime.date(2023, 4, 23), datetime.date(2023, 4, 28)]
Dates in 2023-05:[datetime.date(2023, 5, 2), datetime.date(2023, 5, 2), datetime.date(2023, 5, 4), datetime.date(2023, 5, 2), datetime.date(2023, 5, 4), datetime.date(2023, 5, 14), datetime.date(2023, 5, 17), datetime.date(2023, 5, 15), datetime.date(2023, 5, 16), datetime.date(2023, 5, 23), datetime.date(2023, 5, 1), datetime.date(2023, 5, 22)]
Dates in 2023-06:[datetime.date(2023, 6, 4), datetime.date(2023, 6, 1), datetime.date(2023, 6, 4), datetime.date(2023, 6, 10), datetime.date(2023, 6, 1), datetime.date(2023, 6, 9), datetime.date(2023, 6, 1), datetime.date(2023, 6, 2), datetime.date(2023, 6, 5)]
Dates in 2023-07:[datetime.date(2023, 7, 11), datetime.date(2023, 7, 27)]
Dates in 2023-08:[datetime.date(2023, 8, 29), datetime.date(2023, 8, 21)]
Dates in 2023-09:[datetime.date(2023, 9, 1), datetime.date(2023, 9, 7), datetime.date(2023, 9, 11), datetime.date(2023, 9, 9), datetime.date(2023, 9, 15), datetime.date(2023, 9, 19), datetime.date(2023, 9, 21), datetime.date(2023, 9, 26), datetime.date(2023, 9, 26), datetime.date(2023, 9, 25), datetime.date(2023, 9, 18), datetime.date(2023, 9, 24), datetime.date(2023, 9, 22), datetime.date(2023, 9, 22), datetime.date(2023, 9, 21)]
Dates in 2023-10:[datetime.date(2023, 10, 4), datetime.date(2023, 10, 6), datetime.date(2023, 10, 3), datetime.date(2023, 10, 18), datetime.date(2023, 10, 19), datetime.date(2023, 10, 13), datetime.date(2023, 10, 24), datetime.date(2023, 10, 25), datetime.date(2023, 10, 6), datetime.date(2023, 10, 13), datetime.date(2023, 10, 4)]
Dates in 2023-11:[datetime.date(2023, 11, 21), datetime.date(2023, 11, 11), datetime.date(2023, 11, 29), datetime.date(2023, 11, 28)]
Dates in 2023-12:[datetime.date(2023, 12, 5), datetime.date(2023, 12, 13), datetime.date(2023, 12, 20), datetime.date(2023, 12, 22), datetime.date(2023, 12, 23), datetime.date(2023, 12, 23), datetime.date(2023, 12, 1), datetime.date(2023, 12, 27), datetime.date(2023, 12, 6), datetime.date(2023, 12, 30), datetime.date(2023, 12, 28), datetime.date(2023, 12, 26), datetime.date(2023, 12, 2)]
Dates in 2024-01:[datetime.date(2024, 1, 28), datetime.date(2024, 1, 29), datetime.date(2024, 1, 31), datetime.date(2024, 1, 9)]
Dates in 2024-02:[datetime.date(2024, 2, 23), datetime.date(2024, 2, 26), datetime.date(2024, 2, 21)]
Dates in 2024-03:[datetime.date(2024, 3, 7), datetime.date(2024, 3, 23)]
Dates in 2024-05:[datetime.date(2024, 5, 20), datetime.date(2024, 5, 27), datetime.date(2024, 5, 29)]
Dates in 2024-06:[datetime.date(2024, 6, 12), datetime.date(2024, 6, 14), datetime.date(2024, 6, 16), datetime.date(2024, 6, 19), datetime.date(2024, 6, 11), datetime.date(2024, 6, 19), datetime.date(2024, 6, 24), datetime.date(2024, 6, 20), datetime.date(2024, 6, 11), datetime.date(2024, 6, 14), datetime.date(2024, 6, 30)]
Dates in 2024-07:[datetime.date(2024, 7, 1), datetime.date(2024, 7, 5), datetime.date(2024, 7, 21), datetime.date(2024, 7, 19), datetime.date(2024, 7, 27), datetime.date(2024, 7, 27), datetime.date(2024, 7, 3), datetime.date(2024, 7, 5)]
Dates in 2024-08:[datetime.date(2024, 8, 1), datetime.date(2024, 8, 3), datetime.date(2024, 8, 4), datetime.date(2024, 8, 9), datetime.date(2024, 8, 15), datetime.date(2024, 8, 30), datetime.date(2024, 8, 31), datetime.date(2024, 8, 19), datetime.date(2024, 8, 13)]
Dates in 2024-10:[datetime.date(2024, 10, 7), datetime.date(2024, 10, 8), datetime.date(2024, 10, 6), datetime.date(2024, 10, 2)]
Dates in 2024-09:[datetime.date(2024, 9, 29), datetime.date(2024, 9, 29), datetime.date(2024, 9, 30), datetime.date(2024, 9, 23), datetime.date(2024, 9, 27), datetime.date(2024, 9, 18), datetime.date(2024, 9, 19), datetime.date(2024, 9, 19), datetime.date(2024, 9, 24), datetime.date(2024, 9, 24), datetime.date(2024, 9, 24), datetime.date(2024, 9, 21), datetime.date(2024, 9, 19), datetime.date(2024, 9, 19), datetime.date(2024, 9, 24), datetime.date(2024, 9, 24), datetime.date(2024, 9, 24)]
Dates in 2024-11:[datetime.date(2024, 11, 10), datetime.date(2024, 11, 26), datetime.date(2024, 11, 8), datetime.date(2024, 11, 5), datetime.date(2024, 11, 19), datetime.date(2024, 11, 28), datetime.date(2024, 11, 22), datetime.date(2024, 11, 24)]
Dates in 2024-12:[datetime.date(2024, 12, 5), datetime.date(2024, 12, 16), datetime.date(2024, 12, 16), datetime.date(2024, 12, 8), datetime.date(2024, 12, 2), datetime.date(2024, 12, 4), datetime.date(2024, 12, 7), datetime.date(2024, 12, 7), datetime.date(2024, 12, 1), datetime.date(2024, 12, 28), datetime.date(2024, 12, 9)]
Dates in 2025-01:[datetime.date(2025, 1, 11), datetime.date(2025, 1, 9), datetime.date(2025, 1, 20), datetime.date(2025, 1, 15), datetime.date(2025, 1, 24), datetime.date(2025, 1, 29), datetime.date(2025, 1, 10)]
Dates in 2025-02:[datetime.date(2025, 2, 12), datetime.date(2025, 2, 3), datetime.date(2025, 2, 10), datetime.date(2025, 2, 14), datetime.date(2025, 2, 26), datetime.date(2025, 2, 13), datetime.date(2025, 2, 14), datetime.date(2025, 2, 14), datetime.date(2025, 2, 10), datetime.date(2025, 2, 13)]
Dates in 2025-03:[datetime.date(2025, 3, 11), datetime.date(2025, 3, 1), datetime.date(2025, 3, 2)]
Dates in 2021-11:[datetime.date(2021, 11, 26), datetime.date(2021, 11, 30), datetime.date(2021, 11, 25), datetime.date(2021, 11, 18), datetime.date(2021, 11, 29), datetime.date(2021, 11, 29), datetime.date(2021, 11, 25), datetime.date(2021, 11, 30), datetime.date(2021, 11, 23), datetime.date(2021, 11, 29), datetime.date(2021, 11, 30), datetime.date(2021, 11, 30), datetime.date(2021, 11, 24), datetime.date(2021, 11, 24), datetime.date(2021, 11, 22), datetime.date(2021, 11, 30)]
```

In [11]:

```
# Create DataFrame
df = pd.DataFrame({'date':collection_dates})
df['month_year'] = df['date'].apply(lambda x: x.strftime('%Y-%m'))

df_grouped = df['month_year'].value_counts().sort_index()

# Create line plot
sns.set(style='whitegrid')
plt.figure(figsize=(10,6))
sns.lineplot(x=df_grouped.index, y=df_grouped.values, marker=',', linestyle='-',color='red', linewidth=2, markersize=8)
plt.xlabel('Year-Month', fontsize=14)
plt.ylabel('Number of Sequences Collected', fontsize=14)
plt.title('Frequency of SARS-CoV-2 BA3 collected for each month', fontsize=16)
plt.xticks(rotation=45, ha='right')
plt.tight_layout()
plt.grid(True, linestyle='--', alpha=0.7)
plt.show()
```

BA.3 omicron sub-lineage had a sharp rise when the Omicron variant hot globally in November 2021. However, it was the weakest of the 5 omicron lineages. Though over the years there hasn't been much change in its genetic makeup it did not give rise to sub-variants that caused significant COVID-19 waves. It has been studied less compared to other omicron sub-variants.

However, samples from South Africa collected in October and November 2024 showed that the BA.3 had several amino acids changes mostly in its Spike protein region. These new variants did not seem to cause any different COVID-19 waves but our fear stems from the possibility of becoming a dangerous variant in the future as has been seen from saltation of Omicron which has been the main variant in circulation for more than 2 years.

Of note is the omicron sub-variant Pirola which had saltation as well, the havoc was seen when its progenies started to regain functions and recombining intra-variantly (with other viruses of the same sub-variant designation) and inter-variantly (with other viruses of other sub-variant designation). THe main sub-variants of 2024 have been offspring of this seemingly harmless at first sight Pirola.

Our concern is that saltatory mutations bring unpredicatability of the extent of how these mutations might play out. Notably when they recombine with other sub-variants of SARS-CoV-2.

In [12]:

```
ba3_saltated_spikes = pd.read_csv("saltation patient status.csv")
ba3_saltated_spikes
```

Out[12]:

|  | Virus name | Accession ID | Collection date | Location | Host | Additional location information | Sampling strategy | Gender | Patient age | Patient status | Last vaccinated | Passage | Specimen | Additional host information | Lineage | Clade | AA Substitutions |
| --- | --- | --- | --- | --- | --- | --- | --- | --- | --- | --- | --- | --- | --- | --- | --- | --- | --- |
| 0 | hCoV-19/South Africa/NICD-R00178/2025 | EPI\_ISL\_19771105 | 2025-01-10 | Africa / South Africa / KwaZulu-Natal | Human | NaN | Pneumonia Surveillance | Female | 48 | unknown | NaN | Original | NaN | NaN | BA.3 | GRA | (NSP5\_P132H,Spike\_H69del,NSP3\_G489S,NSP4\_T327I... |
| 1 | hCoV-19/South Africa/NICD-N58822/2024 | EPI\_ISL\_19771107 | 2024-11-22 | Africa / South Africa / Gauteng | Human | NaN | Baseline Surveillance | Male | 5 | unknown | NaN | Original | NaN | NaN | BA.3 | GRA | (NSP5\_P132H,Spike\_H69del,NSP3\_G489S,NSP4\_T327I... |
| 2 | hCoV-19/South Africa/NICD-N58843/2024 | EPI\_ISL\_19771108 | 2024-11-24 | Africa / South Africa / Gauteng | Human | NaN | Baseline Surveillance | Female | 53 | unknown | NaN | Original | NaN | NaN | BA.3 | GRA | (NSP5\_P132H,Spike\_H69del,NSP3\_G489S,NSP4\_T327I... |
| 3 | hCoV-19/USA/CO-CDPHE-43093479/2025 | EPI\_ISL\_19775777 | 2025-02-26 | North America / USA / Colorado | Human | NaN | Baseline surveillance | unknown | unknown | unknown | NaN | Original | NaN | NaN | BA.3 | GR | (NSP5\_P132H,Spike\_H69del,NS7b\_F19L,Spike\_S50L,... |

##### Pairwise Alignment and Number of Mutations in Saltation sequences¶

Jalview was used for Pairwise Alignment between:

> Accession Id: EPI\_ISL\_19771107 (GP1) and EPI\_ISL\_19771108 (GP2) - Isolated in South Africa - Gauteng Province (collectively **GP**)
>
> Accession Id: EPI\_ISL\_19771105 - Isolated in South Africa - KwaZulu Natal (KZN)
>
> Accession id: EPI\_ISL\_402124 - Wuhan Hu 1 Reference sequence (WT)

###### Pairwise Alignements Identities¶

1. GP1 VS GP2 = 100%
2. GP vs KZN = 99.89%
3. GP vs WT = 97.07%
4. WT vs KZN = 97.04%

Of the 3 sequences showing saltation evolution, 2 had identical S gene sequences (from Gauteng Province hence \_gp suffix) collected in Nov 2024, leaving us with 2 different sequences, gp\_sequence and kzn\_sequence (isolated in KwaZulu Natal in Jan 2025).

The amino acid changes positions compared to the Wild type Wuhan Hu 1 Reference are given in the tuples below.

In [13]:

```
aa_mutation_positions_gp = (1162,688,529,244,642,348,704,614,145,147,951,67,654,157,440,251,243,679,339,764,403,796,435,478,371,417,583,445,681,144,9,405,641,137,477,211,136,795,70,452,139,69,373,484,969,655,21,954,138,554,501,460,140,1184,212,852,326,146,143,187,496,142,625,408,101,375,498)
# positions of amino acid poistions unique to Gauteng province compared to Wild type
sorted_gp_mutations = sorted(aa_mutation_positions_gp)  #this puts amino acid changes in order

aa_mutation_positions_kzn = (939,446,575,172,164,141,26,356,688,529,244,642,348,704,614,145,147,951,67,654,157,440,251,243,679,339,764,403,796,435,478,371,417,583,445,681,144,9,405,641,137,477,211,136,795,70,452,139,69,373,484,969,655,21,954,138,554,501,460,140,1184,212,852,326,146,143,187,496,142,625,408,101,375,498)
# amino acid changes positions in KZN sequence compared to Wild Type
sorted_kzn_mutations = sorted(aa_mutation_positions_kzn)

# print(sorted_kzn_mutations)    #testing if the positions are rearranged in ascending order
# Supposed to group these mutations into different domains which are found on the Spike protein. To note where most of the changes are occuring.
```

In [14]:

```
def count_mutations(dataset_name):                    #this function groups, counts and prints number of amino acid changes in each S protein site.

    n_terminal_domain=[]
    receptor_binding_domain = []  
    receptor_binding_motif = [num for num in dataset_name if 437<=num<=501]  # the receptor binding motif is part of the RBD that gets into direct contact with hACE2 receptor
    sd1_sd2 = []     
    s2_subunit = []

    for val in dataset_name:             # the mutation positions are first grouped into their corresponding S protein sites list.
        if val < 318:
            n_terminal_domain.append(val)
        elif 319<=val<=541:
            receptor_binding_domain.append(val)
        elif 531<=val<=591:
            sd1_sd2.append(val)
        else:
            s2_subunit.append(val)


    count_ntd = len(n_terminal_domain)          # the len() counts number of sites in the given list
    count_rbd = len(receptor_binding_domain)
    count_sd1_sd2 =len(sd1_sd2)
    count_s2 = len(s2_subunit)
    count_rbm = len(receptor_binding_motif)

# The next step is to print how many mutations are in each site of the S protein.

    print(f'The N-terminal domain (NTD) which spans from amino acid position ~ 1-318 has {count_ntd} mutations')
    print(f'The receptor binding domain (RBD) which spans from amino acid position ~ 319 - 541 has {count_rbd} mutations with {len(receptor_binding_motif)} mutations specifically in the receptor binding motif ') 
    print(f'The Sub-domain 1 and Sub-domain 2 (SD1/SD2) which span from amino acid position ~ 531 - 591 has {count_sd1_sd2} mutations')
    print(f'The S2 subunit(S2) which spans from amino acid position ~ >592 has {count_s2} mutations')

    sns.set()
    sites = ['NTD','RBD','SD1/SD2','S2']
    frequency=[count_ntd, count_rbd, count_sd1_sd2, count_rbm]
    data = pd.DataFrame({'Sites':sites, 'Frequency of mutations':frequency})
    sns.barplot(x='Sites',y= 'Frequency of mutations', data = data)
    plt.title('Frequency of mutations for every site in dataset')
    plt.show()
```

In [15]:

```
def mutation_plot(dataset_name):                    #this function groups, counts and prints number of amino acid changes in each S protein site.

    n_terminal_domain=[]
    receptor_binding_domain = []  
    receptor_binding_motif = [num for num in dataset_name if 437<=num<=501]  # the receptor binding motif is part of the RBD that gets into direct contact with hACE2 receptor
    sd1_sd2 = []     
    s2_subunit = []

    for val in dataset_name:             # the mutation positions are first grouped into their corresponding S protein sites list.
        if val < 318:
            n_terminal_domain.append(val)
        elif 319<=val<=541:
            receptor_binding_domain.append(val)
        elif 531<=val<=591:
            sd1_sd2.append(val)
        else:
            s2_subunit.append(val)


    count_ntd = len(n_terminal_domain)          # the len() counts number of sites in the given list
    count_rbd = len(receptor_binding_domain)
    count_sd1_sd2 =len(sd1_sd2)
    count_s2 = len(s2_subunit)
    count_rbm = len(receptor_binding_motif)

# The next step is to print how many mutations are in each site of the S protein.

    print(f'The N-terminal domain (NTD) which spans from amino acid position ~ 1-318 has {count_ntd} mutations')
    print(f'The receptor binding domain (RBD) which spans from amino acid position ~ 319 - 541 has {count_rbd} mutations with {len(receptor_binding_motif)} mutations specifically in the receptor binding motif ') 
    print(f'The Sub-domain 1 and Sub-domain 2 (SD1/SD2) which span from amino acid position ~ 531 - 591 has {count_sd1_sd2} mutations')
    print(f'The S2 subunit(S2) which spans from amino acid position ~ >592 has {count_s2} mutations')

     
    sns.set(style='whitegrid', font = 'Arial', font_scale=1.2)

    # Defining the regions and their corresponding amino acid spans
    regions = ['NTD','RBD','SD1/SD2','S2']
    start_positions = [1, 319, 542, 592]
    end_positions = [318, 541, 591, 1273]
    mutations = [count_ntd, count_rbd, count_sd1_sd2, count_s2]

    # Calculation the bin widths and positions
    bin_widths = np.array(end_positions) - np.array(start_positions) + 1
    bin_positions = np.array(start_positions) + np.array(bin_widths) / 2 - 0.5

    #Defining color palette
    colors = sns.color_palette('husl', len(regions))

    # Creating the figure and axis
    fig, ax = plt.subplots(figsize=(10,6))

    # Plotting the histogram with variable-width bins

    for i, (position, width, mutation, color) in enumerate(zip(bin_positions, bin_widths, mutations, colors)):
        ax.bar(position, mutation, width=width, align='center', alpha=0.7, color=color)
    # ax.bar(bin_positions, mutations, width=bin_width, align='center', alpha=0.7)

    # Setting the x-axis ticks and labels
    ax.set_xticks(np.array([(start_positions[i] + end_positions[i])/2 for i in range(len(start_positions))]))
    ax.set_xticklabels(regions, rotation=45, ha='right')

    # Setting the x-axis limits
    ax.set_xlim(0, max(end_positions) + 100)

    # Setting the title and labels 
    ax.set_title("Number of Mutations of SARS-CoV-2 S Protein Region")
    ax.set_xlabel("S Protein Regions")
    ax.set_ylabel('Number of Mutations')

    # Showing the legend
    for i, region in enumerate(regions):
        ax.plot([], [], label=region, color=colors[i], linewidth = 5)
    ax.legend(loc='upper right', bbox_to_anchor=(1.05,1), fontsize=12)

    # Showing the plot
    plt.tight_layout()
    # plt.savefig("mutations_plot.png", dpi=300, bbox_inches='tight')
    plt.show()
```

In [16]:

```
# For KZN sequence
mutation_plot(dataset_name = sorted_kzn_mutations)   #Calling functions to give us the number of mutations in each S protein
                                                        #site followed by a bar graph showing the frequency of mutations for each site

    # both methods can give the same result.
```

```
The N-terminal domain (NTD) which spans from amino acid position ~ 1-318 has 28 mutations
The receptor binding domain (RBD) which spans from amino acid position ~ 319 - 541 has 24 mutations with 11 mutations specifically in the receptor binding motif 
The Sub-domain 1 and Sub-domain 2 (SD1/SD2) which span from amino acid position ~ 531 - 591 has 3 mutations
The S2 subunit(S2) which spans from amino acid position ~ >592 has 19 mutations
```

In [17]:

```
# For GP sequence                                                        
mutation_plot(sorted_gp_mutations)
```

```
The N-terminal domain (NTD) which spans from amino acid position ~ 1-318 has 24 mutations
The receptor binding domain (RBD) which spans from amino acid position ~ 319 - 541 has 22 mutations with 10 mutations specifically in the receptor binding motif 
The Sub-domain 1 and Sub-domain 2 (SD1/SD2) which span from amino acid position ~ 531 - 591 has 2 mutations
The S2 subunit(S2) which spans from amino acid position ~ >592 has 19 mutations
```

The S region mutations in both Gauteng and KwaZulu Natal were fairly higher in the N-terminal domain (NTD), followed by the receptor-binding domain then subunit 2 (S2) with the lowest in subdomain 1 and subdomain 2 intersection (SD1\_SD2). The histograms shown above have different length bins to show coverage of such regions in the S protein. THis helps us to understand there is actually more mutation rates in the RBD than other regions because as it covers less amino-acid regions of S protein it still has several mutations.

##### Phylogenetic Analysis¶

FUBAR automatically removes duplicate sequences to save memory. The Phylogenetic tree information was exported in form of a newick (.new) file which was used in the iTOL.com webserver (interactive Tree of Life) to plot the Phylogenetic tree below.

THe Phylogenetic validates 2 points:

1. The lowest branch represents sequences collected in Nov 2024 and Jan 2025 (labelled in red) which evolved through acquiring many mutations. THe other earlier BA.3 sequences are distantly clustered together (labelled in black) on the Upper branches which validates the new sequences are way distant from earlier BA.3 sequences to an extent even the Wuhan Hu 1 Reference Sequence (Wild Type - Yellow) is closer to each of them than each clusters of BA.3.
2. This information gives us confidence that the calculations of mutation selection rates is very accurate given that its Phylogenetic tree clustered more related sequences together.

##### Calculating Selection Pressure driving evolution in BA.3 lineage¶

Although we have found which sites are mutating, there is also need to know which sites are changing more frequently than the other or which ones are being selected by evolution.

`FUBAR` pervasive selection analysis tool hosted on the `Datamonkey` webserver was used to investigate sites undergoing both positive (changing amino acids) and negative pervasive selection (change in RNA but not the amino acid). Parameters for `FUBAR` analysis were:

> 1. Number of grid points = 20
>
> 1. Concentration parameter of the Dirichlet prior = 0.5
>
> `FUBAR` results were exported in two forms:
>
> 1. Posterior rate distribution exported as an image:

> 1. Table of selection results exported for each site in .csv format shown in the next code cell:

In [19]:

```
fubar_selection_table = pd.read_csv('BA3 datamonkey-table.csv')  
fubar_selection_table.tail()
```

Out[19]:

|  | Site | Partition | alpha | beta | beta-alpha | Prob[alpha>beta] | Prob[alpha<beta] | BayesFactor[alpha<beta] |
| --- | --- | --- | --- | --- | --- | --- | --- | --- |
| 1272 | 1273 | 1 | 1.918 | 1.013 | -0.904 | 0.499 | 0.443 | 0.904 |
| 1273 | 1274 | 1 | 1.635 | 1.154 | -0.481 | 0.487 | 0.454 | 0.948 |
| 1274 | 1275 | 1 | 1.847 | 0.956 | -0.892 | 0.500 | 0.442 | 0.900 |
| 1275 | 1276 | 1 | 1.797 | 10.183 | 8.386 | 0.163 | 0.795 | 4.423 |
| 1276 | 1277 | 1 | 1.541 | 0.995 | -0.546 | 0.491 | 0.450 | 0.931 |

##### Number of sites undergoing pervasive positive selection¶

All sites showing a positive value for `beta - alpha` were recorded as positive sites to understand number of sites undergoing positive selection.

`alpha` = synonymous substitution rate (where a change in a base does not change the final amino acid)

`beta` = non-synonymous substitution rate (where a change in a base changes the final amino acid.)

In [20]:

```
positive_sites = []       # List to hold all sites which are showing positive selcetion

for idx,val in enumerate(fubar_selection_table['beta-alpha']):
    if val > 0:
        positive_sites.append(idx)

    else:
        pass


positive_sites_number = str(len(positive_sites))

print(f'{positive_sites_number} sites show pervasive positive selection')
```

```
71 sites show pervasive positive selection
```

##### Sites undergoing **strong** positive selection¶

The Bayes Factor (BF) from FUBAR results was used to test the hypothesis of pervasive positive selection (β > α) at each site. A higher BF indicates a stronger support for positive selection. Several sites exhibited high BF values, as shown by the peaks in the bar plot below:

In [21]:

```
site = fubar_selection_table['Site']
bayes_factor = fubar_selection_table['BayesFactor[alpha<beta]']

df = pd.DataFrame({'Region' : site, 'Bayes Factor': bayes_factor})


# # Heat map
# bayes_factor_matrix = np.array([bayes_factor])      #   Creating a matrix with 1 row
# plt.figure(figsize=(10,2))
# sns.heatmap(bayes_factor_matrix, cmap='coolwarm', annot=True, xticklabels=site, yticklabels=False)
# plt.xlabel('Site number')
# # plt.ylabel('Bayes Factor')
# plt.title('Bayes factor between sites to show strong positive selection (peaks) in SARS-CoV-2 BA.3')
# plt.show();

# # Scatterplot
# sns.set()
# plt.figure(figsize=(8,6))
# sns.scatterplot(x="Region", y ='Bayes Factor', data=df)
# plt.xlabel('Site number')
# plt.ylabel('Bayes Factor')
# plt.title('Bayes factor between sites to show strong positive selection (peaks) in SARS-CoV-2 BA.3')
# plt.show();


#Barplot
plt.figure(figsize=(16,10), dpi =60)
ax=plt.axes()
ax.plot(site, bayes_factor)
plt.xlabel('Site number')
plt.ylabel('Bayes Factor')
plt.title('Bayes factor between sites to show strong positive selection (peaks) in SARS-CoV-2 BA.3')
plt.show();
```

In [22]:

```
strong_beta = fubar_selection_table['Prob[alpha<beta]']
strong_alpha = fubar_selection_table['Prob[alpha>beta]']

strong_positive_selection = fubar_selection_table.loc[strong_beta > 0.90]
strong_negative_selection = fubar_selection_table.loc[strong_alpha > 0.90]

strong_selection_dfs = [strong_negative_selection,strong_positive_selection]
strong_selection_sites = pd.concat(strong_selection_dfs)
strong_selection_sites
```

Out[22]:

|  | Site | Partition | alpha | beta | beta-alpha | Prob[alpha>beta] | Prob[alpha<beta] | BayesFactor[alpha<beta] |
| --- | --- | --- | --- | --- | --- | --- | --- | --- |
| 67 | 68 | 1 | 31.649 | 1.594 | -30.055 | 0.978 | 0.010 | 0.012 |
| 94 | 95 | 1 | 3.446 | 35.480 | 32.034 | 0.016 | 0.955 | 24.103 |
| 211 | 212 | 1 | 3.422 | 27.351 | 23.929 | 0.036 | 0.930 | 15.076 |
| 342 | 343 | 1 | 3.459 | 25.828 | 22.369 | 0.040 | 0.924 | 13.911 |
| 374 | 375 | 1 | 2.254 | 24.450 | 22.197 | 0.028 | 0.947 | 20.212 |
| 378 | 379 | 1 | 2.099 | 24.596 | 22.498 | 0.026 | 0.951 | 22.049 |
| 408 | 409 | 1 | 3.545 | 25.751 | 22.206 | 0.042 | 0.922 | 13.474 |
| 420 | 421 | 1 | 3.543 | 25.564 | 22.021 | 0.042 | 0.922 | 13.364 |
| 443 | 444 | 1 | 2.763 | 23.087 | 20.324 | 0.038 | 0.931 | 15.322 |
| 449 | 450 | 1 | 3.493 | 28.867 | 25.374 | 0.031 | 0.935 | 16.352 |
| 455 | 456 | 1 | 3.193 | 26.760 | 23.566 | 0.066 | 0.905 | 10.870 |
| 481 | 482 | 1 | 3.461 | 25.602 | 22.141 | 0.041 | 0.924 | 13.748 |
| 487 | 488 | 1 | 3.410 | 30.369 | 26.959 | 0.028 | 0.940 | 17.799 |
| 504 | 505 | 1 | 3.737 | 29.567 | 25.830 | 0.033 | 0.931 | 15.241 |
| 682 | 683 | 1 | 2.763 | 23.087 | 20.324 | 0.038 | 0.931 | 15.322 |
| 684 | 685 | 1 | 3.942 | 44.667 | 40.725 | 0.006 | 0.962 | 29.085 |
| 855 | 856 | 1 | 2.941 | 24.887 | 21.946 | 0.036 | 0.933 | 15.811 |
| 957 | 958 | 1 | 3.927 | 24.311 | 20.384 | 0.051 | 0.909 | 11.334 |

THe table above gives us an idea of the amino acid positions which are undergoing positive selection e.g position 95, 375, 379 and 685 gave us the peaks above BF factor = 20.

##### Heatmap showing which region is undergoing negative or positive selection¶

As we can see, on its own the bar does not give us a summary of which region is undergoing much selection. SO the analysis went further to calculate mean BF values of sites grouped into their corresponding regions.

In [23]:

```
# Defining the regions and their corresponding amino acid spans
region_df = pd.DataFrame({
    'Region': ['NTD','RBD', 'SD1_SD2','S_2'],
    'Start' : [1, 319, 542, 592],
    'End' : [318, 541, 591, 1273]
})

df = pd.DataFrame({
    'site' : fubar_selection_table['Site'],
    'bayes_factor' : fubar_selection_table['BayesFactor[alpha<beta]']
})

# Defining the region positions
region_positions = {
    'NTD': (1, 318),
    'RBD': (319, 541),
    'SD1_SD2': (542, 591),
    'S_2': (592,1270)
}

# Assign each site to a region
def assign_region(site):
    for region, (start, end) in region_positions.items():
        if start <= site <= end:
            return region
    return None

df['Region'] = df['site'].apply(assign_region)

# Calculating the mean Bayes factor for each region
mean_bayes_factor = df.groupby('Region')['bayes_factor'].mean()


print(mean_bayes_factor)

# import seaborn as sns

# import matplotlib.pyplot as plt
 # Creating a dataframe with region names and mean Bayes factors
mean_bayes_factors_df = mean_bayes_factor.reset_index()
mean_bayes_factors_df.columns = ['Regions', 'Mean Bayes Factor']

#Specifying teh order of teh regions
# region_order = ('NTD','RBD','SD1_SD2', 'S_2')

# Pivot the dataframe to create a matrix
mean_bayes_factors_matrix = mean_bayes_factors_df.pivot_table(index=None, columns= 'Regions', values = 'Mean Bayes Factor')
# mean_bayes_factors_matrix = mean_bayes_factors_matrix[region_order]

# Heat map
# bayes_factor_matrix = np.array(mean_bayes_factor)      #   Creating a matrix with 1 row
plt.figure(figsize=(12,3))
sns.heatmap(mean_bayes_factors_matrix, cmap='coolwarm', annot=True, fmt='.3f', linewidths=0.5) #, xticklabels=site, yticklabels=False)
plt.xlabel('S Protein regions', fontsize=14) 
plt.xticks(rotation=45, ha='right')
plt.ylabel('') 
plt.tight_layout()
plt.title('Mean Bayes factor between S protein regions in SARS-CoV-2 BA.3', fontsize=16)
plt.show();
```

```
Region
NTD        1.149528
RBD        1.857691
SD1_SD2    1.092440
S_2        1.111024
Name: bayes_factor, dtype: float64
```

It was seen that most of the selection pressure is in the RBD region. However, it was necessary to investigate further which RBD region is undergoing pervasive selection pressure. THe RBD has a receptor-binding motif(RBM) which directly binds with the human ACE2 receptor, to minimise confusion the RBD was divided into 2 groups the RBD\_A and RBD\_B where the RBM is located.

In [24]:

```
# Defining the regions and their corresponding amino acid spans
region_df = pd.DataFrame({
    'Region': ['NTD','RBD_A','RBM (RBD_B)', 'SD1_SD2','S_2'],
    'Start' : [1, 319, 437, 542, 592],
    'End' : [318, 436, 501, 591, 1273]
})

df = pd.DataFrame({
    'site' : fubar_selection_table['Site'],
    'bayes_factor' : fubar_selection_table['BayesFactor[alpha<beta]']
})

# Defining the region positions
region_positions = {
    'NTD': (1, 318),
    'RBD_A': (319, 436),
    'RBM (RBD_B)': (437,501),
    'SD1_SD2': (542, 591),
    'S_2': (592,1270)
}

# Assign each site to a region
def assign_region(site):
    for region, (start, end) in region_positions.items():
        if start <= site <= end:
            return region
    return None

df['Region'] = df['site'].apply(assign_region)

# Calculating the mean Bayes factor for each region
mean_bayes_factor = df.groupby('Region')['bayes_factor'].mean()


print(mean_bayes_factor)

# import seaborn as sns

# import matplotlib.pyplot as plt
 # Creating a dataframe with region names and mean Bayes factors
mean_bayes_factors_df = mean_bayes_factor.reset_index()
mean_bayes_factors_df.columns = ['Regions', 'Mean Bayes Factor']

#Specifying teh order of teh regions
# region_order = ('NTD','RBD','SD1_SD2', 'S_2')

# Pivot the dataframe to create a matrix
mean_bayes_factors_matrix = mean_bayes_factors_df.pivot_table(index=None, columns= 'Regions', values = 'Mean Bayes Factor')
# mean_bayes_factors_matrix = mean_bayes_factors_matrix[region_order]

# Heat map
# bayes_factor_matrix = np.array(mean_bayes_factor)      #   Creating a matrix with 1 row
plt.figure(figsize=(12,3))
sns.heatmap(mean_bayes_factors_matrix, cmap='coolwarm', annot=True, fmt='.3f', linewidths=0.5) #, xticklabels=site, yticklabels=False)
plt.xlabel('S Protein regions', fontsize=14) 
plt.xticks(rotation=45, ha='right')
plt.ylabel('') 
plt.tight_layout()
plt.title('Mean Bayes factor between S protein regions in SARS-CoV-2 BA.3', fontsize=16)
plt.show();
```

```
Region
NTD            1.149528
RBD_A          1.727517
RBM (RBD_B)    2.295831
SD1_SD2        1.092440
S_2            1.111024
Name: bayes_factor, dtype: float64
```

Most of the selection pressure is in the RBM, if there are more mutation rates its not long until it adapts quickly to bind efficiently to the hACE2. However, when designing and assigning vaccines and/or drugs it is wiser to target regions undergoing less selection rates or more conserved regions. Apparently, the NTD-directed vaccines and drugs can work in treating BA.3. Although NTD is not the most conserved region, it is the most viable choice due to its well-researched target for therapeutics.

In [ ]:
